# Non-Negative Data-Driven Mapping of Structural Connections in the Neonatal Brain

**DOI:** 10.1101/2020.03.09.965079

**Authors:** E. Thompson, A.R. Mohammadi-Nejad, E.C. Robinson, M.F. Glasser, S. Jbabdi, M. Bastiani, S.N. Sotiropoulos

**Affiliations:** Sir Peter Mansfield Imaging Centre, School of Medicine, University of Nottingham, Nottingham, United Kingdom; Nottingham National Institute of Health Research (NIHR) Biomedical Research Centre, Queen’s Medical Centre, University of Nottingham, United Kingdom; School of Biomedical Engineering and Imaging Sciences, King’s College London, London, United Kingdom; Department of Neuroscience, Washington University School of Medicine, Saint Louis, USA; Department of Radiology, Washington University School of Medicine, Saint Louis, USA; Wellcome Centre for Integrative Neuroimaging - FMRIB, University of Oxford, Oxford, United Kingdom

## Abstract

Mapping connections in the neonatal brain can provide insight into the crucial early stages of neurodevelopment that shape brain organisation and lay the foundations for cognition and behaviour. Diffusion MRI and tractography provide unique opportunities for such explorations, through estimation of white matter bundles and brain connectivity. Atlas-based tractography protocols, i.e. a priori defined sets of masks and logical operations in a template space, have been commonly used in the adult brain to drive such explorations. However, rapid growth and maturation of the brain during early development make it challenging to ensure correspondence and validity of such atlas-based tractography approaches in the developing brain. An alternative can be provided by data-driven methods, which do not depend on predefined regions of interest. Here, we develop a novel data-driven framework to extract white matter bundles and their associated grey matter networks from neonatal tractography data, based on non-negative matrix factorisation that is inherently suited to the non-negative nature of structural connectivity data. We also develop a non-negative dual regression framework to map group-level components to individual subjects. Using in-silico simulations, we evaluate the accuracy of our approach in extracting connectivity components and compare with an alternative data-driven method, independent component analysis. We apply non-negative matrix factorisation to whole-brain connectivity obtained from publicly available datasets from the Developing Human Connectome Project, yielding grey matter components and their corresponding white matter bundles. We assess the validity and interpretability of these components against traditional tractography results and grey matter networks obtained from resting-state fMRI in the same subjects. We subsequently use them to generate a parcellation of the neonatal cortex using data from 323 new-born babies and we assess the robustness and reproducibility of this connectivity-driven parcellation.

## Introduction

The neonatal period is a critical time for brain development, during which the refinement and maturation of white matter connections lay the groundwork for later cognitive development (Ball et al., 2015; Counsell et al., 2008; Girault et al., 2019). With diffusion MRI (dMRI) we can track these connections non-invasively and *in vivo*, which enables us to study the early development of structural connectivity and microstructure, even during the first weeks of life (see (Ouyang et al., 2019) for a recent review).

DMRI studies of neonates have shown that the trajectory of fibre maturation reflects the neurodevelopmental hierarchy, with primary motor and sensory tracts developing earlier than the association tracts that enable higher order functioning (Dubois et al., 2008; Kulikova et al., 2015; Partridge et al., 2004). Studies have also demonstrated the impact of preterm birth (Ball et al., 2015; Batalle et al., 2017b; Brown et al., 2014; Girault et al., 2019) and maternal environment (Deoni et al., 2013; Tam et al., 2016) on the early development of white matter.

Despite the large potential of diffusion imaging for exploring early developmental stages of the brain, current analysis techniques follow the paradigms that have been established for the adult brain. For instance, dMRI tractography protocols for identifying specific white matter bundles typically rely on delineation of regions of interest (ROIs) that provide a priori anatomical knowledge on the route of the tract; and these ROIs can be defined relative to a template for automated delineation (Bastiani et al., 2019; de Groot et al., 2013; Warrington et al., 2019).

However, neonatal brains are not simply small adult brains (Batalle et al., 2017a), and this renders the above paradigm problematic. The rapid growth and changes in brain morphology during the neonatal period, as well as fast alterations in tissue composition that alter imaging contrast over time (Bastiani et al., 2019), render it challenging to ensure correspondence between template-driven ROIs and tractography protocols at different stages of development (Serag et al., 2012). Manual delineation on a subject-by-subject basis could be an alternative, but it is time-consuming, assumes very detailed knowledge of how neonatal neuroanatomy is depicted in MRI at various early development stages, and becomes prohibitive for large cohorts, such as the developing human connectome projects (Howell et al., 2019; Hughes et al., 2017).

In this paper, we propose an alternative approach for simultaneously mapping white matter bundles and the corresponding grey matter nodes in the neonatal brain using data-driven methods, which are inherently model-free and are expected to be more immune to the challenges described above. Independent component analysis (ICA) has been a commonly used data-driven method for identifying brain networks from resting-state functional MRI (fMRI) data (McKeown et al., 1998), and recent work has shown that it can be also applied to dMRI tractography data of the adult human brain (O’Muircheartaigh and Jbabdi, 2017; Wu et al., 2015) or of the non-human primate brain (Mars et al., 2019). We develop an alternative approach to ICA and explore its applicability in the neonatal brain.

One limitation of applying ICA to tractography data is that the estimated independent components and the respective mixing matrix can contain both positive and negative values, whereas structural connectivity data is inherently non-negative. This leads to challenges in the interpretation of negative weights. To address this problem, we present an alternative data-driven method that can be used to identify non-negative connectivity components. Our approach is based on non-negative matrix factorisation (NMF) (Lee and Seung, 2001). Like ICA, NMF is an unsupervised technique that estimates a pre-defined number of components from the data. However, the elements and their weights are constrained to take non-negative values. Sparsity constraints in the decomposition allow identifiability and further provide an indirect means of requiring independence between the estimated components. This results in a set of components whose weighted summation represents the whole system. Due to these advantageous properties, NMF has been recently used to identify networks of structural covariance (Ball et al., 2019; Sotiras et al., 2017, 2015) from MRI data.

In this study, we present for the first time an NMF-based framework for extracting connectivity components from diffusion MRI data, both at the group and the individual level. We apply this approach within the context of mapping patterns of structural connections in new-born babies, aged 37 to 44 weeks post-menstrual age (PMA) at scan, using publicly-released data provided by the developing Human Connectome Project (dHCP) (Hughes et al., 2017; Hutter et al., 2018). First, we describe the theory for decomposing whole-brain tractography-induced connectivity matrices into grey matter networks and their corresponding white matter bundles. We subsequently use simulations to quantitatively evaluate the behaviour of the method and assess its performance against ICA. We explore the validity and interpretability of i) the automatically detected white matter patterns against results from standard tractography protocols available through the dHCP (Bastiani et al., 2019) and ii) the grey matter patterns against components obtained from data-driven mapping of resting-state fMRI in the same subjects. Finally, we use the extracted structural connectivity components from a group of 323 new-born babies to derive connectivity-driven cortical parcellations of the neonatal brain and assess their robustness and reproducibility.

## Theory

Let **X** be an *M x N* dense^1^ “connectivity” matrix, with X_ij_ {*i=1:M, j=1:N}* carrying information on the likelihood of structural connections existing between locations *i* and *j* in the brain. Without loss of generality, let us assume that locations *i* represent the whole brain and comprise of all imaging voxels, and that locations *j* represent grey matter and reside on the cortical white/grey matter boundary (WGB) and in subcortical grey matter. Diffusion MRI tractography can provide such a matrix if we seed streamlines from *N* seeds on the WGB and subcortical nuclei, and record visitation counts to *M* voxels across the brain, such that each column of **X** describes the connectivity profile of a grey matter location *j*. A data-driven decomposition of **X** can identify *K* components based on similarity of connectivity profiles. Different numbers of components can be obtained depending on the desired properties of the estimated components.

**Independent component analysis** (ICA) imposes statistical independence between the components to perform a linear decomposition. An observed matrix **X** is represented as **X** = **WS**, where **S** is the independent sources matrix (each row *k* corresponds to a source/ component) and **W** the weights or mixing matrix (each column *k* corresponds to the weights of source *k*). As this is an ill-posed problem in general, ICA uses source independence to estimate an un-mixing matrix **A**, that best approximates **W**^-1^, to recover the original sources from the observed data: **AX** ≈ **S**. This process is entirely data-driven by the statistical properties of the mixture, with no prior knowledge of the mixing matrix or the signals. The first step of all ICA algorithms is to centre and whiten the data, to remove linear dependencies from the data. This can be achieved with a principal component analysis (PCA) or singular value decomposition (SVD). Then we seek an orthogonal rotation **V** to apply to the whitened data to optimise the statistical independence of the estimated components. This cannot be done analytically but there are a number of different methods of solving the problem iteratively. The FastICA algorithm (Hyvärinen and Oja, 2000), which uses non-Gaussianity as a proxy for independence, is one of the typically used algorithms.

ICA has been used to identify networks from resting-state functional MRI data (McKeown et al., 1998), where *T* is the number of timepoints and the decomposition results into *K* spatial maps (covering all *N* brain voxels), each with a weight vector of length *T.* Each weight *w*_*ik*_ represents how much component *k* contributes to activity recorded at time point *i*. ICA has also been used recently in the case of dMRI tractography, where *N* is the number of seeds (O’Muircheartaigh and Jbabdi, 2017). In that case, the decomposition provides *K* spatial maps (covering all *N* points on the grey matter), each representing a component with shared connectivity profile through white matter, associated with a weight vector of length *M*. Each weight *w*_*ik*_ represents in this case how much component *k* contributes to the connection patterns of voxel *i*.

**Non-negative matrix factorisation** is an alternative decomposition technique, where a matrix **X** is factorised into two matrices **W** and **H**, under the constraint that all three contain only positive values (Lee and Seung, 1999). This is more naturally suited for use with structural connectivity data, which is inherently non-negative. In general, NMF is an ill-posed problem and there exist multiple solutions in most cases. The linear superposition of components, combined with the non-negativity constraint, lead to an implicit sparsity constraint in the algorithm (requesting a signal to be explained as a linear combination of non-negative regressors will inherently lead many weights close to zero). Additional explicit sparsity constraints can be applied to further constrain the solution space and improve the identifiability of the decomposition (Hoyer, 2004). Specifically, the cost function *C* to minimise is of the form: 

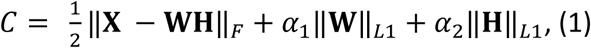

where ‖*x*‖_*F*_ is the Frobenius norm, ‖*x*‖_*L*1_ is the *L*_*1*_-norm, used to explicitly impose sparsity, and α_1_ and α_2_ are tuning parameters that allow us to control the degree of regularisation on the mixing matrix and component matrix, respectively. Higher values of α’s lead to more sparsity in the resultant decomposition. The NMF can be initialised with a non-negative SVD, which has been shown to improve the accuracy of the decompositions (Boutsidis and Gallopoulos, 2008).

### Dimensionality reduction and back-projection

In certain cases, large *M* dimensions (i.e. large number of imaging voxels) can pose computational and numerical convergence challenges. One can therefore use PCA to reduce the *M x N* matrix **X**, into a *P x N* matrix **X**_**r**_ of principal components. Applying the decomposition to this reduced matrix, results in a *K x N* set of components **S**, and a *P x K* mixing matrix in PCA space **W**_**r**_. In order to obtain the mixing matrix in the original space of *M* imaging voxels, we can take the pseudoinverse of the component matrix **S** and project it back onto the original data to obtain the tract space mixing matrix, i.e. **W = XS**^**†**^, where **S**^**†**^ denotes the pseudoinverse of **S** (see Suppl. Figure 1).

### From group to subject decompositions - Non-negative dual regression

When considering data matrices **X** from multiple subjects (e.g., by averaging across subjects in the simplest case), the components and mixing matrices will represent the group. Dual regression can then be used to generate subject-level representations of the group components and mixing matrices (Beckmann et al., 2009; Nickerson et al., 2017), both for ICA and NMF decompositions. Dual regression comprises of two steps:

i. Identify the subject-specific mixing matrix 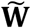 from the group-level grey matter components **S**, using the subject-level connectivity matrix 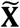 

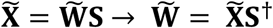
ii. Find the subject-level grey matter components 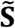, using the subject-specific mixing matrix 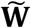: 

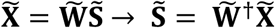

In previous work, this multivariate regression has been achieved by taking the pseudo-inverse of the group-level components and the subject-level mixing matrix (O’Muircheartaigh and Jbabdi, 2017), as illustrated in Suppl. Figure 2. However, taking the pseudoinverse introduces negative values into the components and their weights, which leads to mixed-sign subject-level representations of the original non-negative group-level components. Instead, we have developed a “non-negative dual regression” technique for back projecting NMF results, using non-negative least squares (NNLS) (Ling et al., 1977) for the regression steps. NNLS solves an equation of the form *argmin*_**x**_||**Ax** − **y**||_2_ subject to **x** ≥0, in which **x** and **y** are vectors, and **A** is a matrix. Thus, the optimisation has to be performed separately for each target voxel in step (i) and each grey matter seed in step (ii) (see Suppl. Figure 3). This provides an entirely non-negative framework for dual regression that retains the sparse characteristics of the group-level NMF components, as shown in figure 2.

**Figure 1.**
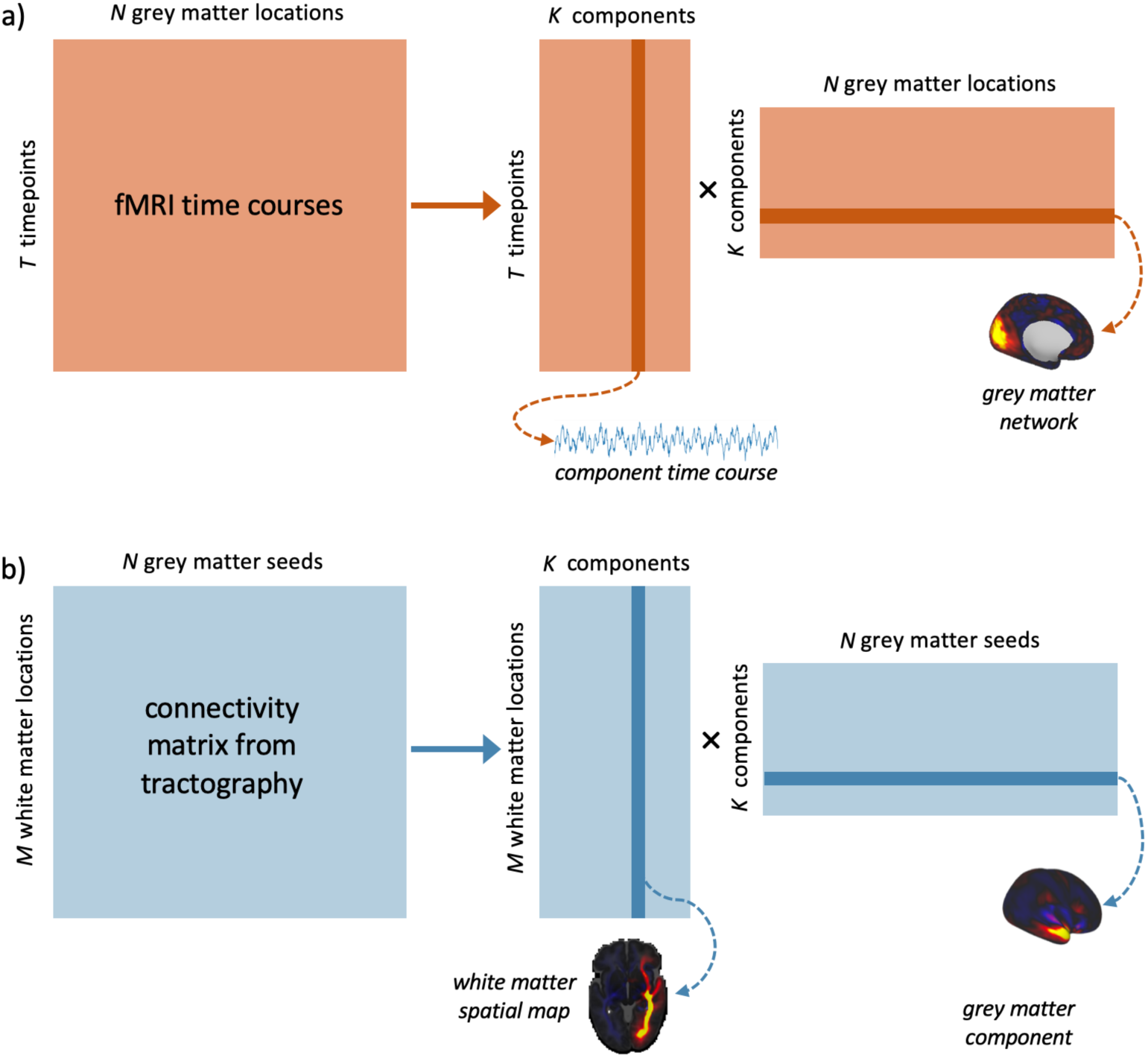
Data-driven matrix decomposition methods applied to resting-state functional MRI and structural connectivity data. a) N functional time-courses of length T are recorded from points in the grey matter. We can apply a matrix decomposition technique, such as ICA, to this matrix, yielding an TxK mixing matrix of time courses and a KxN matrix of spatial components. b) an MxN connectivity matrix describes the likelihood of structural connections existing between each of N grey matter seeds and M locations in the brain. The equivalent decomposition applied to this matrix gives us an MxK mixing matrix of spatial maps, and a KxN matrix of components in the grey matter.

**Figure 2.**
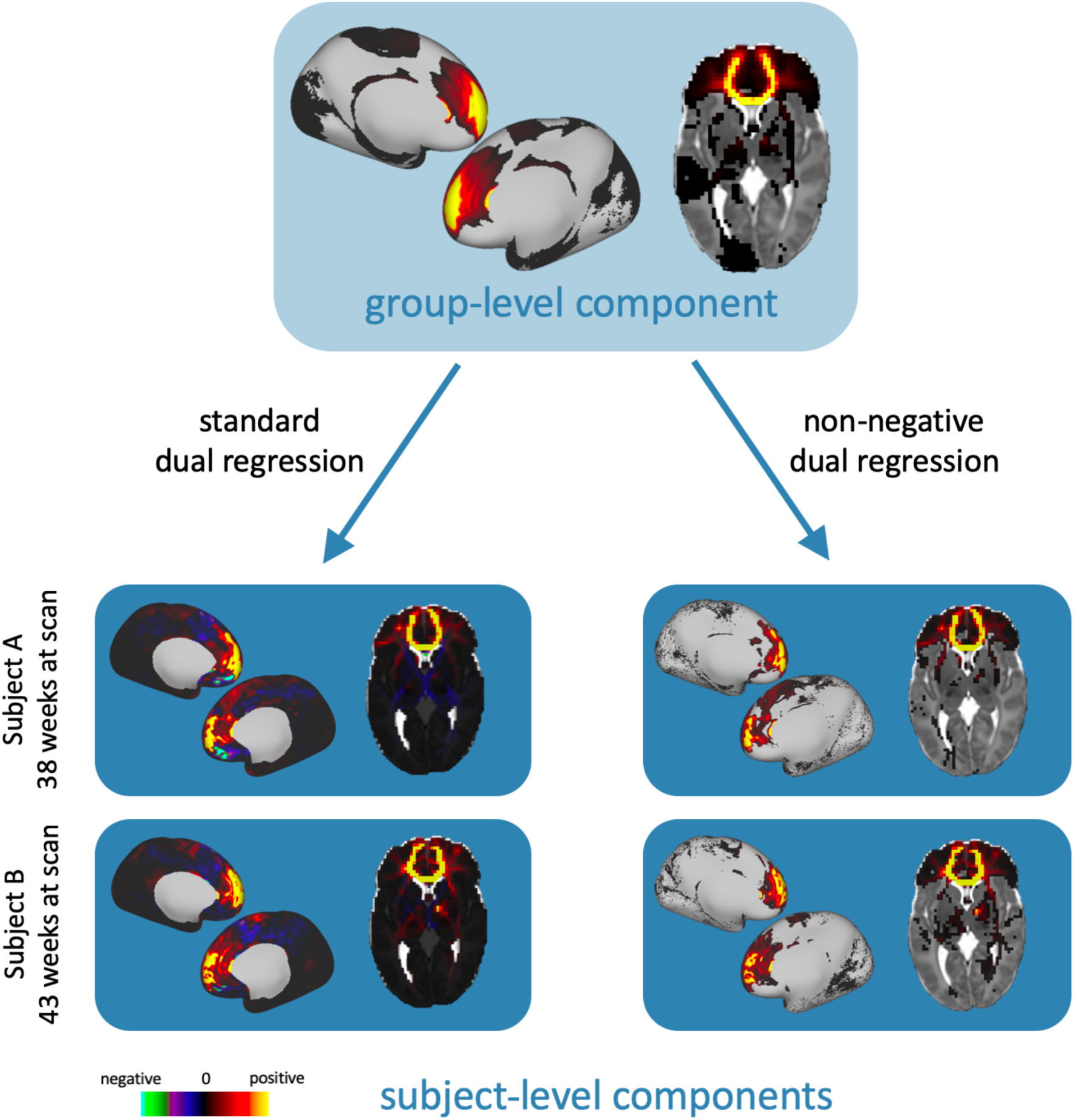
Example dual regression results for a component from a K = 50 NMF decomposition. On the left, the component has been dual regressed onto two subjects’ data with the standard approach using the pseudoinverse. On the right, the component has been dual regressed with our non-negative method that uses non-negative least squares. In all volumetric images, left is left.

## Methods

We present the development of the above frameworks to map structural connectivity in the neonatal brain. We first give an overview of the data employed. We then describe a set of simulations that allow principled evaluation of the decomposition frameworks and we finally describe the methods we use to illustrate the benefits of our approach.

### Data

We used structural, dMRI, and fMRI data made publicly available by the developing Human Connectome Project (dHCP) (www.developingconnectome.org). Briefly, data were acquired during natural sleep on a 3T Philips Achieva with a dedicated neonatal imaging system, including a neonatal 32 channel head coil (Hughes et al., 2017; Hutter et al., 2018). Diffusion MRI data were acquired over a spherically optimised set of directions on three shells (b = 400, 1000 and 2600 s/mm^2^, total number of volumes acquired per subject: 300). Pre-processing was carried out according to the dHCP diffusion processing pipeline (Bastiani et al., 2019). This includes motion correction and distortion correction (Andersson et al., 2016; Andersson and Sotiropoulos, 2016). The data were super-resolved along the slice direction to achieve isotropic resolution of 1.5 mm^3^. Cortical surface reconstruction was carried out from T2w images with an isotropic resolution of 0.5 mm^3^, using a pipeline specifically adapted for neonatal structural MRI data (Makropoulos et al., 2018). Resting-state functional MRI data were acquired for 15 minutes (TE/TR = 38/392 ms, 2300 volumes) with an acquired resolution of 2.15 mm isotropic. fMRI pre-processing was carried out as detailed in (Fitzgibbon et al., 2019), with an automated pipeline including fieldmap pre-processing to estimate susceptibility distortion; registration steps; susceptibility and motion correction; and denoising with ICA-FIX.

Data were considered from a group of 323 subjects born at term age (175 male, 148 female). Median (range) birth age was 40.1 (37.0, 42.3) postmenstrual weeks and age at scan40.9 (37.4, 44.4) weeks. Pre-processed data are available through the latest dHCP’s data release^2^.

### Data processing and whole-brain tractography

Pre-processed data were further analysed to obtain structural connectivity matrices. To ensure alignment between subjects, we registered the anatomical surfaces to a representative template space before performing tractography. First, we used a surface registration pipeline (https://github.com/ecr05/dHCP_template_alignment), based on the multi-modal surface matching (MSM) algorithm (Robinson et al., 2018, 2014). Cortical folding was used to drive the alignment of neonatal WGB, cortical mid-thickness, and pial surfaces to the dHCP 40-week PMA surface templates (Bozek et al., 2018). This aligned vertices on the WGB surface to ensure consistent seed points for tractography across subjects. We then applied a previously computed non-linear volumetric registration (ANTs, Avants et al., 2011) to all MSM-derived surfaces to register them to 40-week PMA volumetric template space (Serag et al., 2012). This step was necessary to ensure that the tractography seeds were aligned to the target space, because the volumetric and surface-based neonatal templates are not inherently aligned (Bozek et al., 2018; Serag et al., 2012).

Once the surfaces were aligned, we obtained connectivity matrices **X** for each subject, by performing whole-brain probabilistic tractography using FSL (Behrens et al., 2007; Hernandez-Fernandez et al., 2019). Fibre orientations (up to 3 per voxel) were estimated using a model-based deconvolution against a zeppelin response kernel, to accommodate for the low anisotropy inherent in data from this age group (Bastiani et al., 2019; Hernández et al., 2013; Sotiropoulos et al., 2016). We subsequently seeded 10,000 streamlines from each of 58,551 vertices on the WGB of both hemispheres (average vertex spacing 1.2 mm, excluding the medial wall) and from each of 2548 subcortical 2mm voxels (bilateral amygdala, caudate, hippocampus, putamen and thalamus), giving us a total of *N* = 61,099 seeds. This type of grey matter seeding has been shown to suffer less from the gyral bias in tractography, compared to whole-brain white matter seeding, even if gyral bias is less prominent in the neonatal brain (Thompson et al., 2019). Visitation counts were recorded between each seed point and each of *M* = 50,272 voxels in a whole-brain mask with the ventricles removed, down-sampled to 2 mm_3_. The pial surface was used as a termination mask to prevent streamlines from crossing between gyri, and streamlines were not allowed to cross the WGB more than twice (once at the seed point and again at termination), to reduce false positives. All masks (seeds, targets, exclusions) were defined in 40 post-menstrual weeks volume template space (Serag et al., 2012), however tractography was carried out in native space with results resampled directly to template space. Visitation counts were multiplied by the length of the pathway to correct for compound uncertainty in the estimated trajectories (O’Muircheartaigh and Jbabdi, 2017). The resulting dense matrices describe the likelihood of a white matter connection between each grey matter seed and the rest of the brain. The connectivity matrices were normalised by the total number of viable streamlines before being averaged across the group.

### Simulations

We evaluated the performance of the decomposition frameworks (using NMF and ICA) in numerically simulated data, before applying them to real data. We simulated datasets **X** with a known number of underlying sources **S**, to observe how the behaviour of the decompositions over different model orders reflects the true dimensionality of the data. To find a realistic generative distribution to use for our sources, we used the spatial maps from standard tractography protocols in the neonatal brain (Bastiani et al., 2019) to generate connectivity blueprints (Mars et al., 2018) as proxies for the source spatial maps in grey matter space (Figure 1), and fit several distributions to the intensities of these maps (unwrapped to 1D). We found that log-beta distributions best described the data. The sources were therefore drawn from log-beta distributions, whose parameters in turn were drawn from Gaussian distributions according to the fits to the measured data. These sources are random and sparse, features that indirectly ensure a high degree of independence. Sources were scaled to lie in the range 0-1. The mixing matrix was randomly generated, normalised so the columns sum squared to 1. The simulated data was calculated as the product of the mixing matrix with the source matrix. Zero-mean, additive Gaussian noise was applied to that product via a logit transform, to maintain non-negativity.

#### Varying L_1_-norm regularisation in NMF

The NMF decomposition can be regularised with *L*_*1*_-norm terms to promote sparsity in the components (see equation (1)) (Févotte and Idier, 2011). We first tested NMF on the simulated data with varying levels of regularisation to assess its effect on the accuracy and robustness of the decomposition. Data were simulated with *K* = 50 sources, and overall dimensions of *N* = 1200 and *M* = 1000, with noise added with σ^2^ = 0.05 to best match the real data. We used the same regularisation parameter for the mixing matrix and the components, i.e. α_1_=α_2_=α, following the implementation in scikit learn (Pedregosa et al., 2011). NMF was applied with model orders from 1 to 100 and with regularisation parameters, α = 0, 0.1, 0.25, 0.5. This process was repeated with 100 noisy realisations of the data in each case.

#### Varying number of sources

We performed the simulations with varying number of sources in the data to check how this affects the results. The data were generated with σ^2^= 0.05 and with *K*=25, 50 and 75 sources. ICA and NMF were applied with model orders from 1 to 100. For NMF, we used a regularisation parameter of α = 0.1 (see Simulation Results for justification). This was also repeated 100 times. ICA was first initialised with a PCA with *P* = 100 components, as described in the theory section.

#### Varying noise levels

Finally, we tested the impact of varying noise levels on the decompositions. Data were simulated as above. Gaussian noise was added to the data with varying σ^2^ = 0.0005, 0.005, 0.05, and 0.5. 100 noisy realisations were generated in each case. The data were decomposed with ICA and NMF, with model orders *K* from 1 to 100. ICA was first initialised with a PCA with *P* = 100 components, as above.

#### Assessing Performance

We used three different metrics to assess the success of the decompositions on the simulated data: i) *Reconstruction error:* the sum of squared errors between the reconstructed data after decomposition and the original data: i.e. Σ(**X** – **WS**)^2^. This gives us a measure of the information lost through the decomposition. ii) *Source-component correlation:* the correlation between each original source and the estimated components. The best-matched component to each source was identified and the mean of the maximum correlation values for each component was considered. This describes how well the decompositions have characterised the underlying signals in the data, and is sensitive to overfitting, as redundant components that are not well matched to sources will bring the value down. iii) *Sparsity:* Following the approach in (Hoyer, 2004; Sotiras et al., 2015), we used a sparsity measure for the derived components based on the relationship between the *L*_*1*_-norm and the *L*_*2*_-norm: 

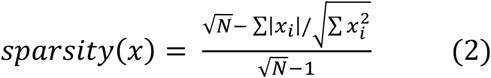

This returns values between 0 and 1, with 1 signifying a maximally sparse component with only one non-zero element. This was calculated for each component vector in **S**, and we report the mean value across all components. Sparse components are desirable because they provide an easily interpretable representation of the data with minimal redundant information. In the case of NMF, sparsity constraints also make results more reproducible, by constraining the solution space. For ICA, sparsity can be thought of as a proxy for independence.

### In-vivo data decompositions

For real data, we decomposed group-average tractography matrices, using independent component analysis (ICA) and non-negative matrix factorisation (NMF), with a range of model orders *K*. ICA was initialised with regular PCA, in which the first 500 components were retained (explaining 97% of the total variance). ICA was applied to the reduced dataset using the FastICA algorithm (Hyvärinen and Oja, 2000), with independence imposed in the seed domain. The pseudo-inverse of this matrix was projected back onto the group-level connectivity matrix to yield the corresponding white matter components. To deal with the sign ambiguity of ICA, components that were negative in the long tail of their distribution were sign-flipped, for consistency with the other methods (i.e. so that the main mass of the distribution was in the positive valued domain).

NMF was performed with a coordinate descent algorithm (Cichocki et al., 2006), a Frobenius norm cost function (see equation (1)), and an *L*_*1*_-norm regularisation parameter α = 0.1. In NMF, the matrix is decomposed directly into the *M x K* mixing matrix and the *K x N* component matrix so there is no need for the back-projection step that was carried out for ICA after the PCA.

All decompositions were implemented using scikit learn (Pedregosa et al., 2011) (python 2.7) and the code is available on GitHub (https://github.com/ethompson93/Data-driven-tractography).

### Comparison to tractography-derived white matter tracts

To assess validity and interpretability of the extracted components, we compared the automatically extracted white matter components with results obtained from standard, template-driven tractography protocols, developed for neonatal subjects, as described in (Bastiani et al., 2019). 28 tracts (13 bilateral) were mapped in each subject. The tracts included in this analysis were: acoustic radiation (AR), anterior thalamic radiation (ATR), cingulate gyrus part of cingulum (CGC), parahippocampal part of cingulum (CGH), cortico-spinal tract (CST), forceps minor (FMI), forceps major (FMA), fornix (FOR), inferior fronto-occipital fasciculus (IFO), inferior longitudinal fasciculus (ILF), medial lemniscus (ML), posterior thalamic radiation (PTR), superior longitudinal fasciculus (SLF), superior thalamic radiation (STR), and uncinate fasciculus (UNC). These were registered to a 40-week template and down sampled to 2 mm for comparison with the tract-space representations of our data-driven components.

### Split-half reliability analysis

We performed a split-half analysis on a cohort of 323 term-age subjects to see how robust and reproducible our decompositions were across different model orders. We evaluated a number of model orders: *K* = 5, 10, 25, 50, 100, 20. For each value of *K*, we performed a one-to-one matching of components across the split-half, based on the Pearson’s correlation coefficients of their spatial maps, recording the correlation coefficients of the matched pairs as a measure of their similarity. This was repeated for the grey matter and white matter maps. We also measured the reconstruction error and the sparsity of the components for both ICA and NMF, as in the simulations.

### Comparison to functional resting-state networks

The cortical patterns of structural connectivity from our NMF components were compared with resting state networks from fMRI. For this analysis, we selected a group of 55 subjects all born and scanned between 40 weeks and 41 weeks PMA (ie. all subjects within this age range who had both structural and functional data available).

We first mapped the functional data onto the cortical surface, broadly following the fMRISurface pipeline outlined in (Glasser et al., 2013). The native WGB, midthickness and pial surfaces were affine registered to the same space as the functional data. The fMRI timeseries were then mapped onto the cortical surface using a partial volume weighted ribbon-constrained volume to surface mapping algorithm, as implemented in HCP’s connectome workbench (Marcus et al., 2011). These data were then downsampled from the native mesh and registered to the 32k resolution template (using the same MSM transform as for the WGB surface used to seed tractography). Spatial smoothing was applied over the cortical surface with a Gaussian kernel, FWHM = 2 mm.

Temporally-concatenated group-ICA was performed using FSL’s Melodic (Beckmann and Smith, 2004), with MIGP for the PCA step (Smith et al., 2014). We specified 50 independent components. We performed NMF on the group-averaged structural connectivity matrices of the same group of subjects, with *K* = 50, for comparison. The similarity between the resultant grey matter spatial maps was assessed using Pearson’s correlation coefficient.

### Cortical parcellations using structural connection patterns

We used the grey matter components to generate a hard parcellation of the cortex with *K* clusters, using a “winner-takes-all” approach. Each vertex on the cortical surface was labelled according to the component that had the highest weighting at that point. We tested the robustness of these parcellations by calculating the Dice coefficient between parcellations generated on each of the split-halves. Dice measures the overlap between two clusters, normalised by the number of elements in each cluster.

We also assessed the parcellation using a Silhouette coefficient, which assesses the similarity of the vertices in a cluster in relation to the vertices in other clusters (Rousseeuw, 1987). We used (1 - Pearson’s R) as a distance metric for the connectivity profiles of different vertices. A successful parcellation would group vertices with similar connectivity profiles, which are distinct from the connections in other parcels.

Results from our data-driven parcellations were benchmarked against a “null distribution” of 100 random Voronoi parcellations of the same model order (Aurenhammer, 1991). Voronoi parcellations are spatially-contiguous and geodesic-distance based and were generated from seeds randomly distributed over the surface of two spheres, mapped to the surface of each hemisphere of the cortex. Each vertex on the sphere is labelled according to its nearest seed point on the surface. The spherical parcellations were projected onto the cortex, providing random parcellations with a set number of contiguous spatial regions.

## Results

### Simulations

We performed simulations to evaluate the performance of ICA and NMF decompositions on a synthetic dataset in which the underlying sources were known. We first looked at the effect of varying the degree of *L*_*1*_-norm regularisation in NMF. We then investigated how the number of sources and the noise level in the data affected results.

#### Varying L_1_-norm regularisation

Increasing the regularisation parameter, α, increases sparsity, but also increases the reconstruction error. The NMF decomposition breaks down for high regularisation (α = 0.5), with high error and very low source-component correlation. Smaller amounts of regularisation improve the agreement between the components and sources and reduce the reconstruction error at the cost of reducing sparsity. A good middle-ground solution is shown (α = 0.1), balancing reconstruction accuracy and sparsity. We therefore opted to use α = 0.1 for subsequent experiments.

#### Varying number of sources

We carried out the decompositions on data with varying numbers of underlying sources. Figure 4 shows that reconstruction error increases with the number of sources, so more information is lost between the decomposition and the original data as the data become more complex. For the source-component correlation, we can see two different regimes. When the number of components, *N*, is lower than the true number of sources in the data, *K*, the average correlation between the components and the true sources rises quickly for very low *N*, then plateaus until *N* = *K*. When *N* > *K*, the extra components overfit to the noise and bring down the average correlation with the sources. NMF achieves overall very high correlations between the reconstructed components and the true non-negative sources. NMF component sparsity increases rapidly for low *N*, then increases more slowly once the number of components exceeds the number of sources. In the case of ICA, sparsity reaches a peak when the number of components is equal to the number of underlying sources, then decreases.

**Figure 3.**
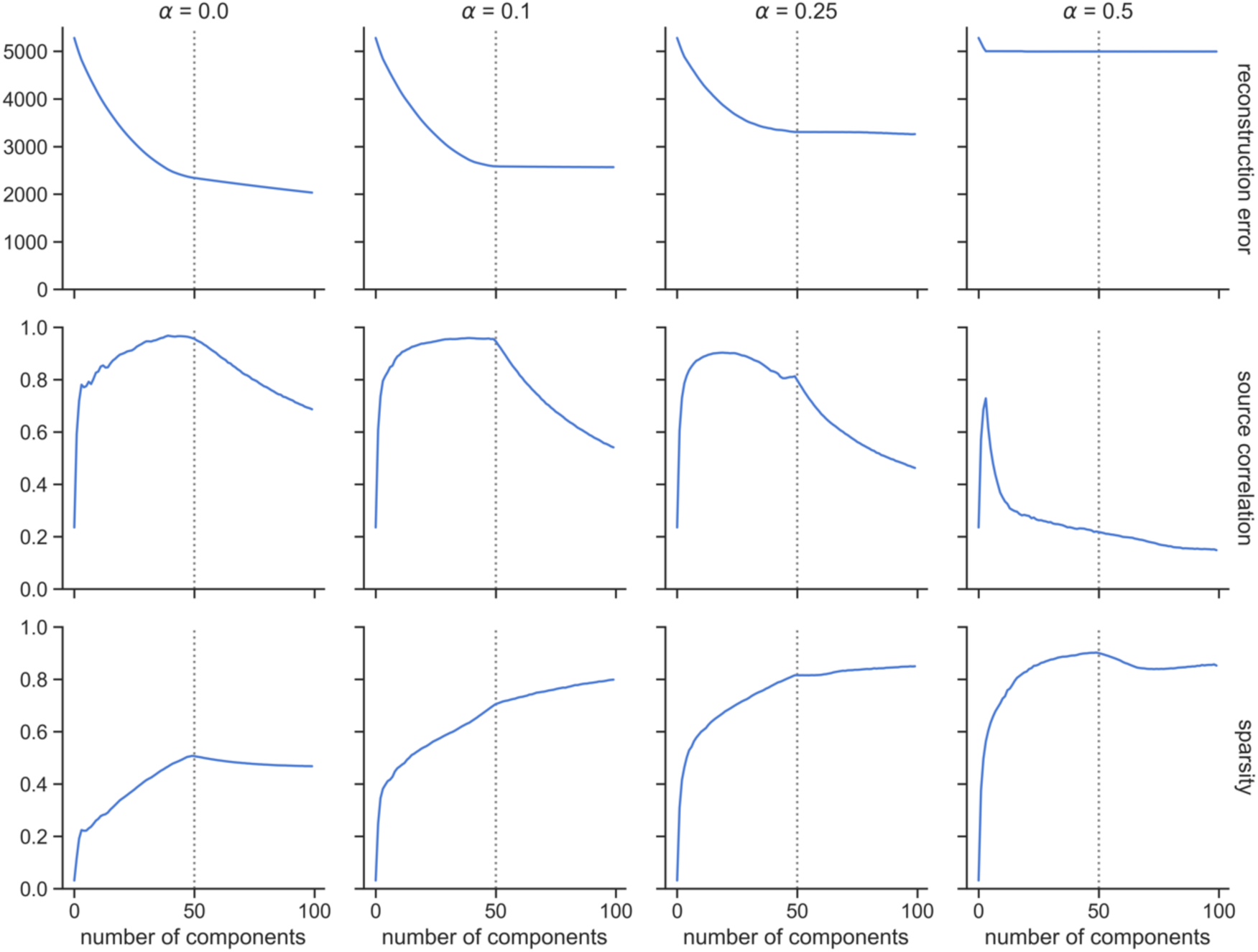
Simulation experiment to assess the effect of L_1_-norm regularisation on NMF. The degree of regularisation increases from left to right across the plots (a = 0.0, 0.1, 0.25, 0.5). Different metrics are shown from top to bottom: reconstruction error, correlation between derived components and underlying sources, sparsity of components. The true number of underlying sources (K = 50) is denoted by a vertical dashed line. Noise variance was σ^2^ = 0.05. Results are shown averaged over 100 noisy realisations of the data.

**Figure 4.**
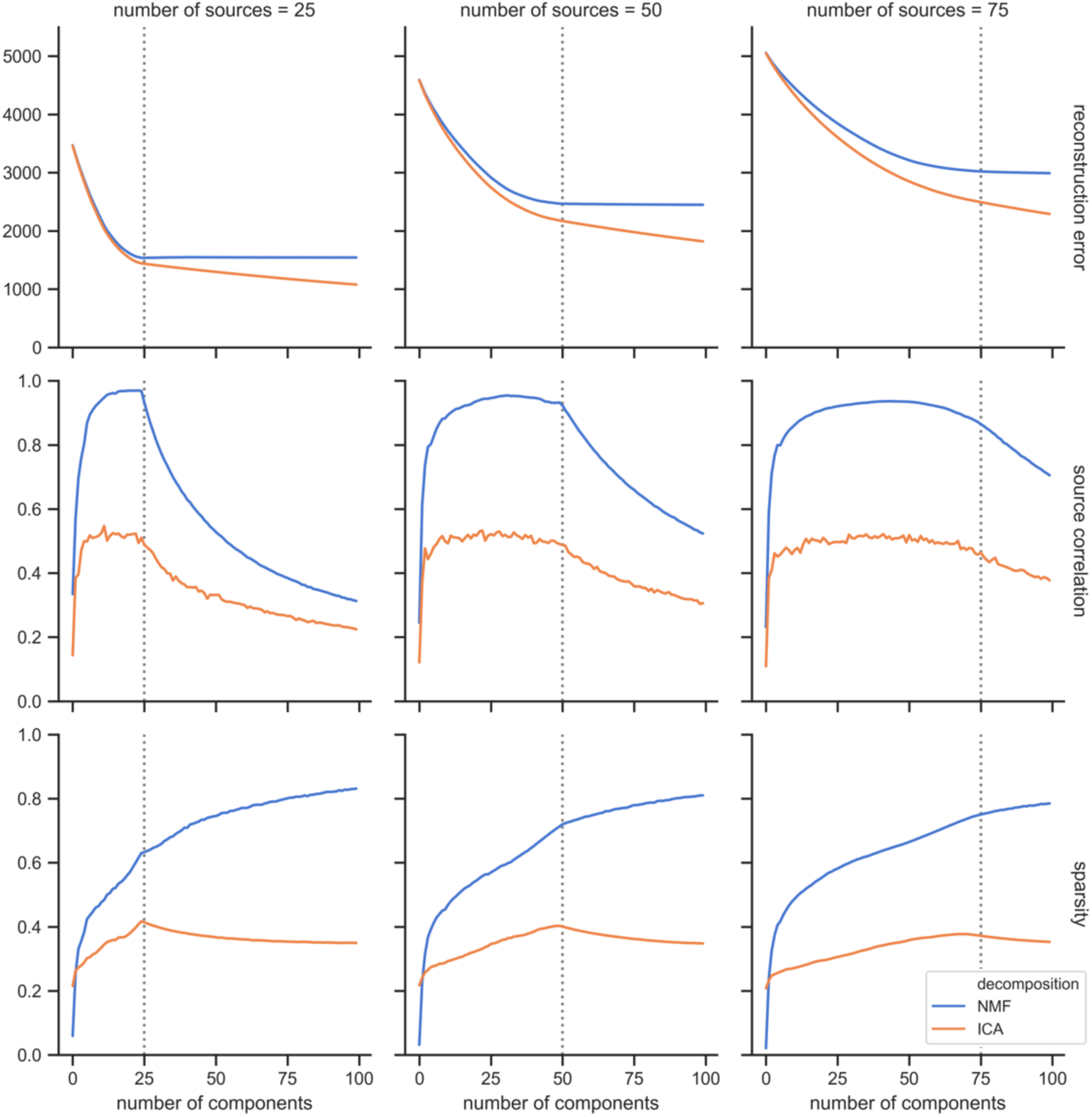
Simulation results to show how decompositions vary with differing numbers of underlying sources. The dotted vertical line shows the number of underlying sources in each case (from left to right: K = 25, 50, 75). Results are shown from ICA and NMF decompositions, in orange and blue, respectively. σ^2^ = 0.05 and a = 0.1 for NMF.

#### Varying SNR

Overall, reconstruction error increases with noise level. In general, reconstruction error decreases as the model order approaches *K*, the true number of underlying sources and then plateaus for higher model orders. The mean correlation between the components and the underlying non-negative sources increases as the number of components approaches *K*, and then decreases as the models overfit to noise. The sparsity of the components exhibits a relatively stable pattern for low and mid-levels of noise, but it becomes considerably reduced in the high noise scenario (σ^2^ = 0.5).

Figures 4 and 5 also enable us to compare the performances of ICA and NMF on simulated, non-negative data. ICA shows a lower reconstruction error than NMF, particularly when model order exceeds the number of true sources. This could, however, signify that ICA is overfitting to noise more than NMF, particularly since ICA also exhibits a lower correlation between its components and the underlying sources than NMF, at all model orders. This reflects the better suitability of NMF for identifying inherently non-negative patterns within the data, in contrast to ICA, which generates components that contain both positive and negative values. NMF also generates components with consistently higher sparsity than those from ICA.

**Figure 5.**
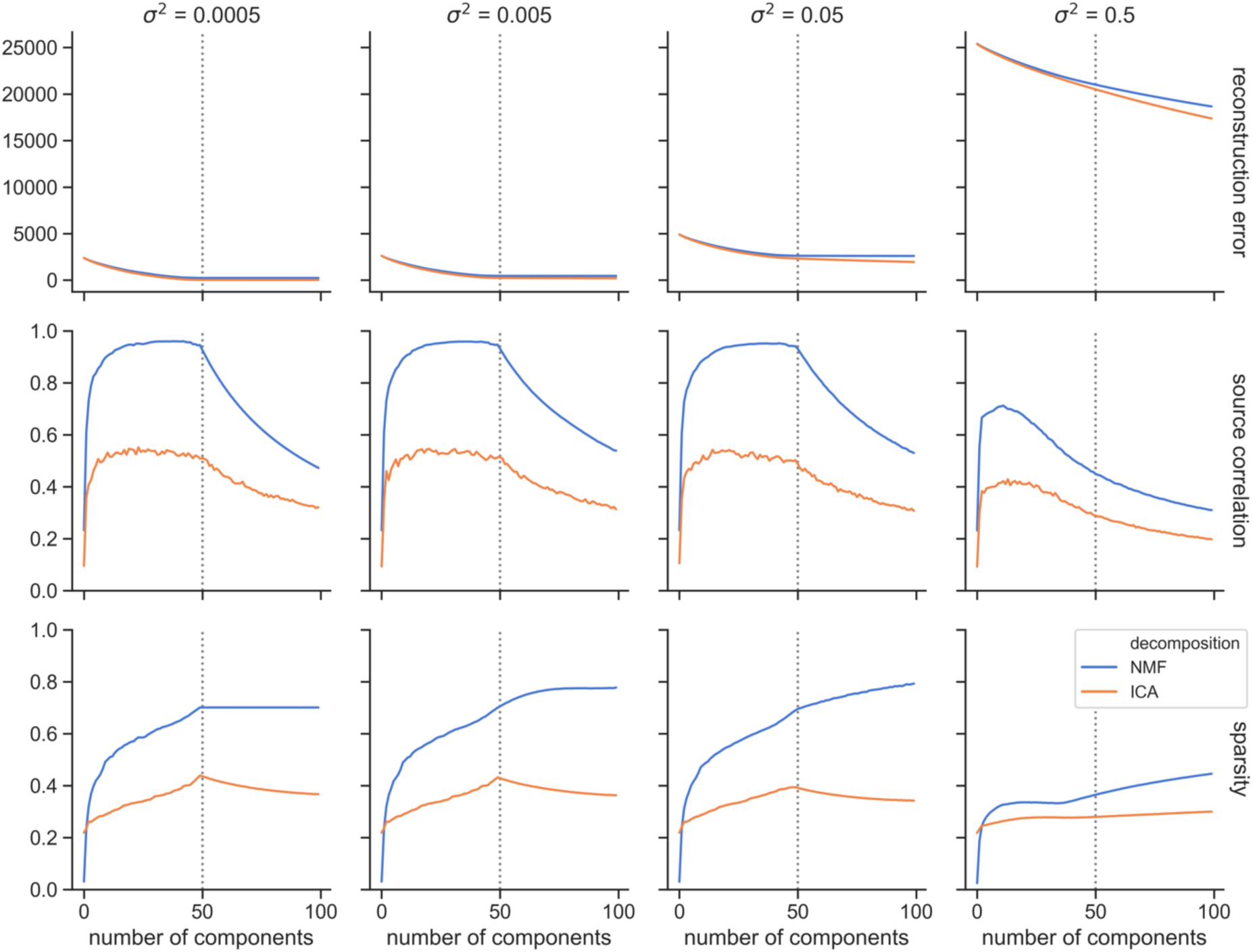
Simulation results to assess the effect of varying noise levels on the ICA (orange) and NMF (blue) decompositions. The noise level increases from left to right across the plots (σ^2^ = 0.0005, 0.005, 0.05, 0.5). Different metrics are shown from top to bottom: reconstruction error, correlation between derived components and underlying sources, sparsity of components. The true number of underlying sources (K = 50) is denoted by a vertical dashed line.

To summarise, we evaluated the performance of ICA and NMF on a simulated dataset with non-negative sources. Based on the results of these simulations, we have chosen a regularisation parameter of α = 0.1 for NMF to use on the real data, as this promotes sparsity in the components, without compromising too much accuracy in the reconstruction. We have found that NMF has a number of advantages over ICA for non-negative data: it generates components that are more closely matched to the real sources, with higher sparsity and potentially less overfitting to noise.

### In-vivo results - Comparison between ICA, NMF and standard tractography

To investigate the interpretability and validity of the extracted components, we compared the white matter components from both ICA and NMF with the group-averaged results from standard tractography protocols. A number of our data-driven components exhibit strong spatial similarity to known white matter pathways (figure 6). In fact, all the considered 28 tracts have well-matching components (Suppl. Figure 4). Both ICA and NMF are able to identify spatially separate regions of grey matter (i.e. networks), along with their underlying white matter connections, for example in the forceps minor, the ILF and the various thalamic projections.

**Figure 6.**
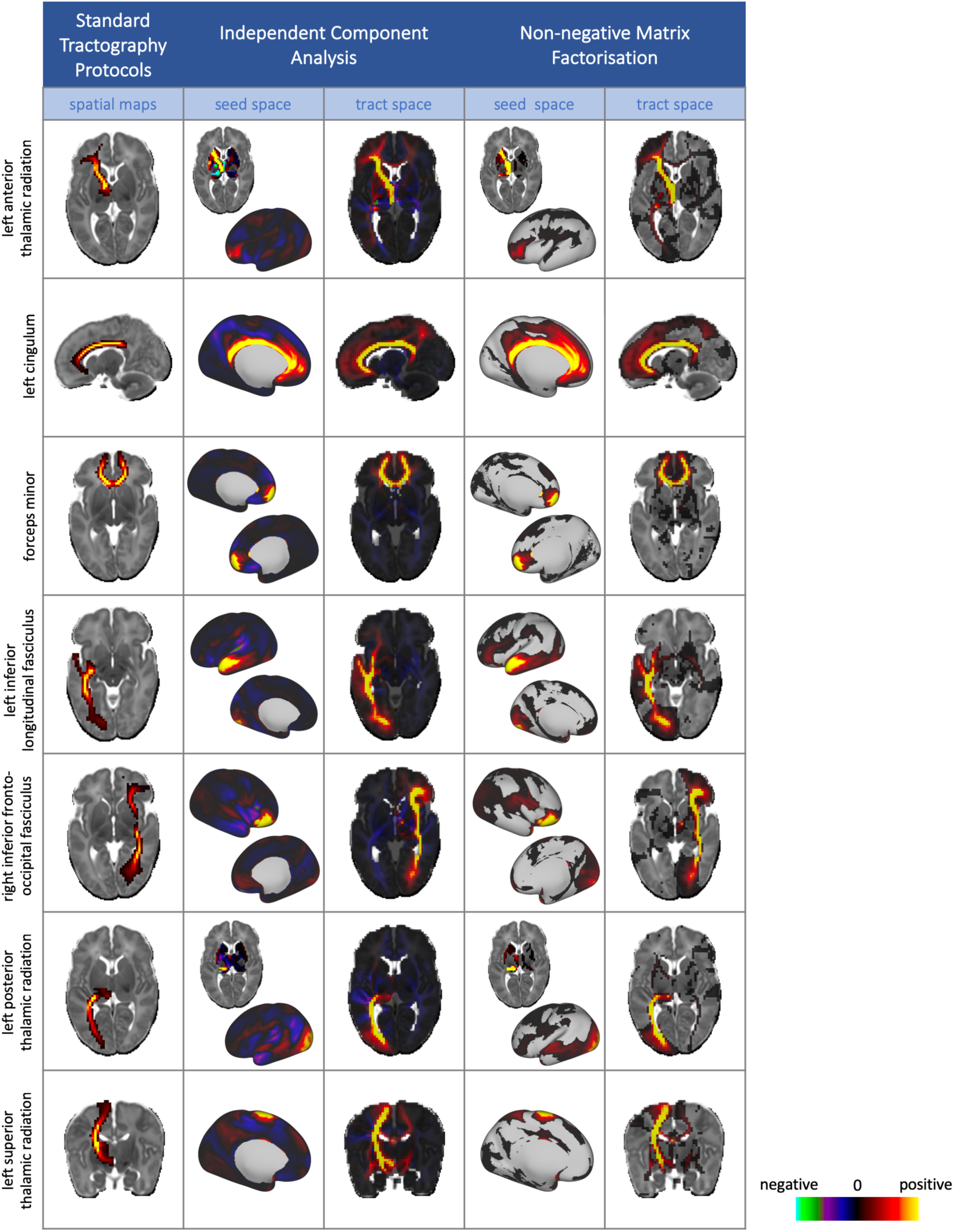
Example group level results from NMF and ICA (model order = 100), displayed alongside their matching tract from the standard protocols. Data-driven components are un-thresholded to enable the comparison between the negative values in the ICA components and the sparse, non-negative representations from NMF. All tractography and data-driven results are taken from split 1 of the split-half analysis. The full set of 28 tracts with their matched data-driven components are shown in the supplementary material.

These examples demonstrate the advantages of using NMF over ICA. NMF components are inherently sparser (ICA-derived spatial maps typically cover the whole brain) and by construction non-negative. The main body of the anatomically relevant information conveyed by ICA components is present with NMF decompositions but in an inherently non-negative manner. This suggests that the NMF sparsity constraints effectively enforce independence in the composition, similarly to ICA. In addition, we can observe qualitative improvements of NMF over ICA for a number of tracts. For instance, the NMF component corresponding to the right IFO has a stronger peak in the occipital lobe than the equivalent ICA component, and NMF has fewer false positive frontal projections in the left ILF.

Interpretability can be also illustrated for components that do not match any tracts from the set we reconstructed using standard tractography protocols. An example is demonstrated in figure 7, where 10 components from the *K =* 100 NMF decomposition have been identified as corresponding to different segments of the corpus callosum. For each component, the grey matter (seed space) map is shown, along with the WM spatial map (tract space).

**Figure 7.**
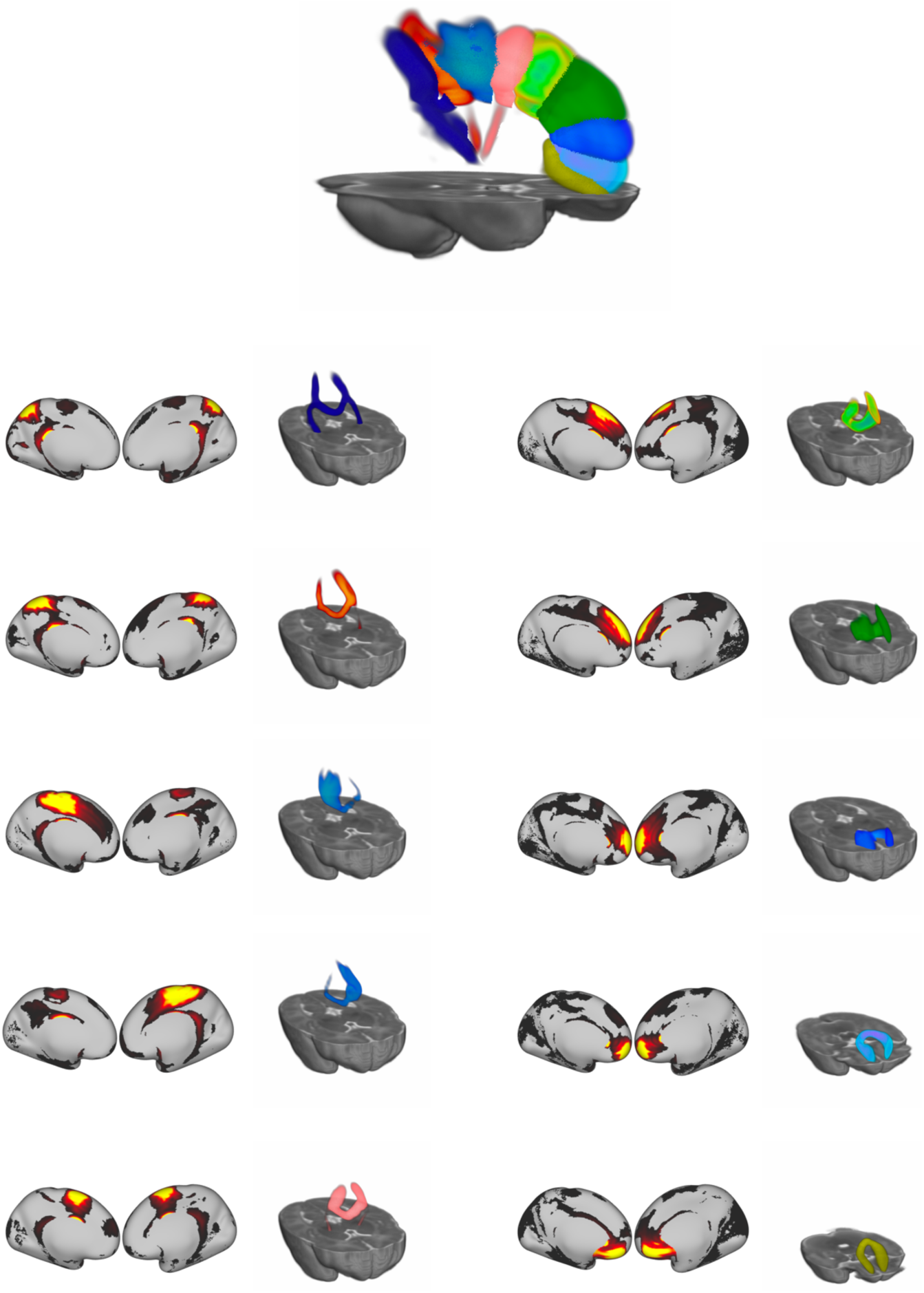
Ten components from the K = 100 NMF decomposition that correspond to segments of the corpus callosum. For each component, the grey matter (seed space) map is shown, along with the WM spatial map (tract space) rendered in 3D to aid visualisation. All rendered WM segments are shown at the top.

### Assessing the reliability and accuracy of the decompositions

To assess the reproducibility of the derived components, we performed a split-half reliability analysis for the ICA and NMF decompositions. Figure 8 presents histograms of correlations between the best-matching components across the split-halves, for both ICA and NMF. In all cases, the median value lies above 0.8, which shows that both methods are robust to different subject groups. Even if patterns are more variable for lower model orders (*K* < 25), both methods perform similarly for higher *K* (50, 100, 200). Similar behaviour is observed for grey matter components and white matter mixing matrices.

**Figure 8.**
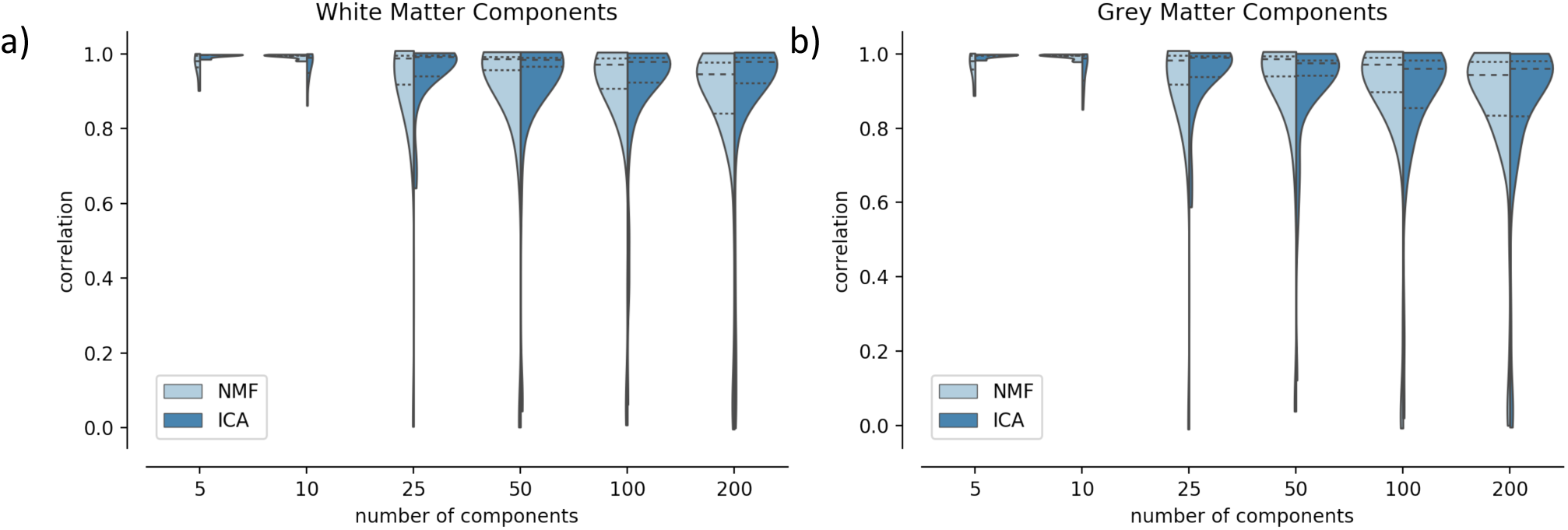
Split-half reliability analysis for ICA and NMF. Pearson’s correlation scores were calculated between the best-matched components in each split for the white matter spatial maps (a) and the grey matter maps (b). The dotted lines on the violin plots indicate the 25^th^ and 75^th^ percentiles and the median is represented by a dashed line.

We also computed the reconstruction error and component sparsity. In line with the results from the simulations, reconstruction error decreases with increasing numbers of components, with ICA having slightly higher reconstruction accuracy than NMF (Suppl. Figure 5). Sparsity is much higher for NMF than for ICA, as we would expect from a qualitative examination of the components in figure 6. Sparsity increases rapidly from 5 to 50 components and increases after 100 components become smaller. Both measures have been calculated for both splits, and confidence intervals are displayed but very small, which indicates that these measures are stable for different groups of subjects.

We explored how increasing the model order in the decomposition affects the splitting of components (Suppl. Figure 6). Equivalent components were identified across model orders by calculating the correlations between their spatial maps. We can see that the more coarse-grained connectivity patterns from the low dimensionality decompositions are broken down into more sparse, fine-grained spatial maps as we increase the number of components. For example, in the left panel of Suppl. Figure 6, we show an NMF component and the associated white matter spatial map from the *K* = 5 decomposition that delineates the left pyramidal tract. As we increase the number of components from *K* = 5 to *K* = 50, we see this bundle split into sub-components that characterise different parts of corona radiata projections. We can also see the increase in sparsity between the low and the high order components (which agrees with the quantitative results - Suppl. Figure 5b).

Having ascertained the reliability of the data-driven framework for a large group of subjects, we explored the behaviour of smaller groups. We performed a *K* = 50 decomposition on a single subject’s data, and then for groups of 5, 10, 50 and 200 subjects. The white matter and grey matter spatial maps from two of the resultant components are shown in figure 9. This shows that the patterns are robust even at the single-subject level, although the patterns are noisier with fewer subjects. A quantitative analysis of the similarity between the small group-size results and the full cohort components is shown in Suppl. Figure 7, from which we can see that components from 10 subjects and 50 subjects have similarly very strong correspondence with the full cohort, while even the single-subject results are reasonable.

**Figure 9.**
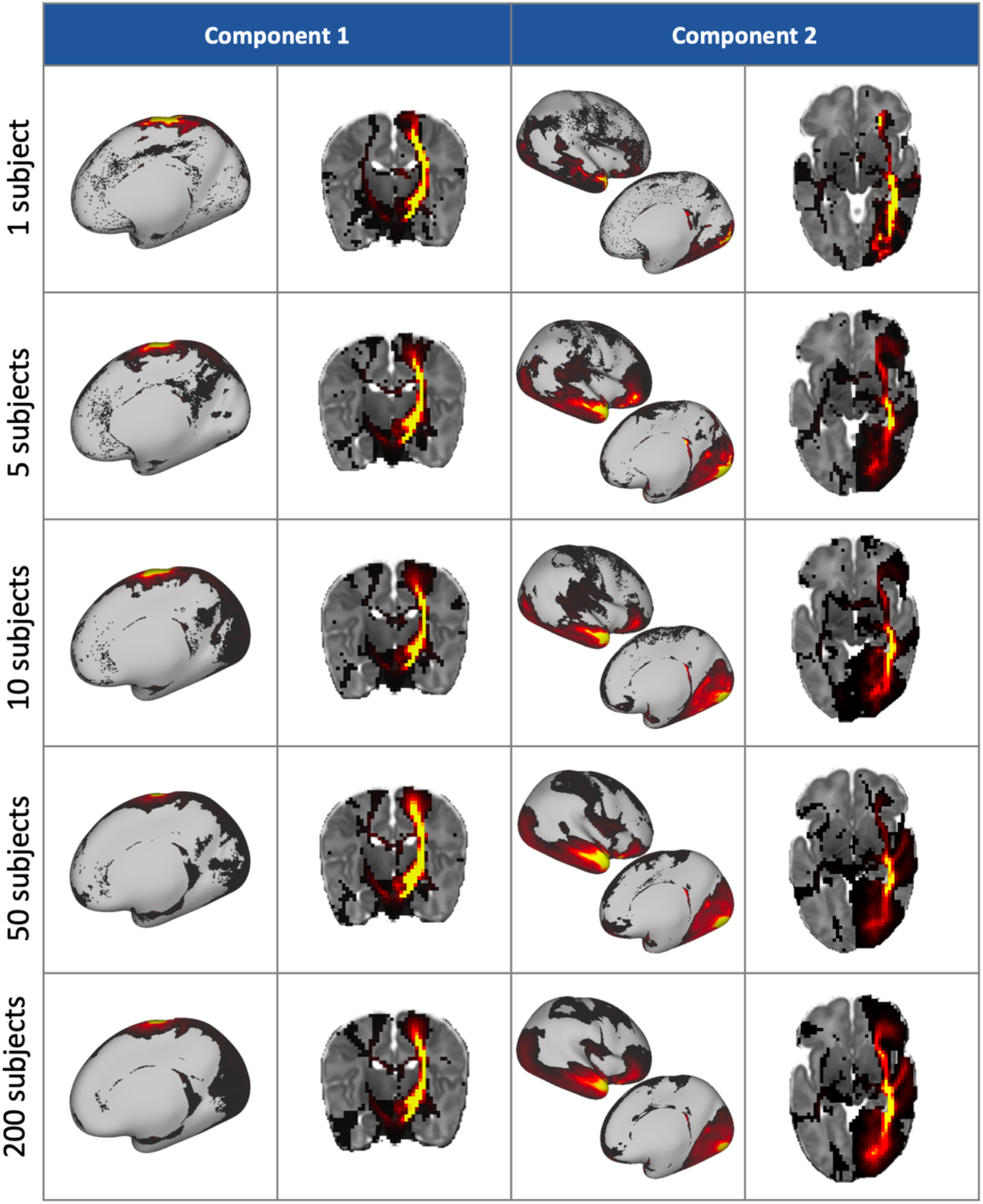
Two components and their corresponding white matter pathways from K = 50 group-level decompositions with varying numbers of subjects. Component 1 correlates well with the tractography-delineated cortico-spinal tract, and component 2 with the inferior longitudinal fasciculus.

### Comparison with functional resting-state networks

As an extra indirect validation, we compared the grey matter maps from the NMF decompositions of the tractography data, with resting-state networks (RSNs) obtained from ICA decomposition of fMRI data. We performed group-level ICA (*K* = 50) on fMRI data from 55 subjects and compared the resultant resting-state networks to those from a *K* = 50 NMF decomposition of the structural connectivity data from the same subjects. Through visual inspection, 24 of the functional components were found to contain noise or artefacts, so were discarded. We measured the similarity of the remaining 26 RSNs to our structural grey matter components using Pearson’s correlation coefficient, r, to identify the best matching pairs.

Most functional components were well matched to at least one structural component, with the lowest correlation value between an RSN and a tractography component being r = 0.2. Over half (14 out of the 26 networks identified) had a correlation value r > 0.5 with their best-matched structural component. The correlation matrix in figure 10 is sparse, which indicates that there is specificity in the matching. Where RSNs were strongly associated with multiple structural components, this was either a bilateral network split into the two hemispheres (e.g. fig 14b and c) or structural networks that overlapped with different regions of the RSN (fig 14a and d).

**Figure 10.**
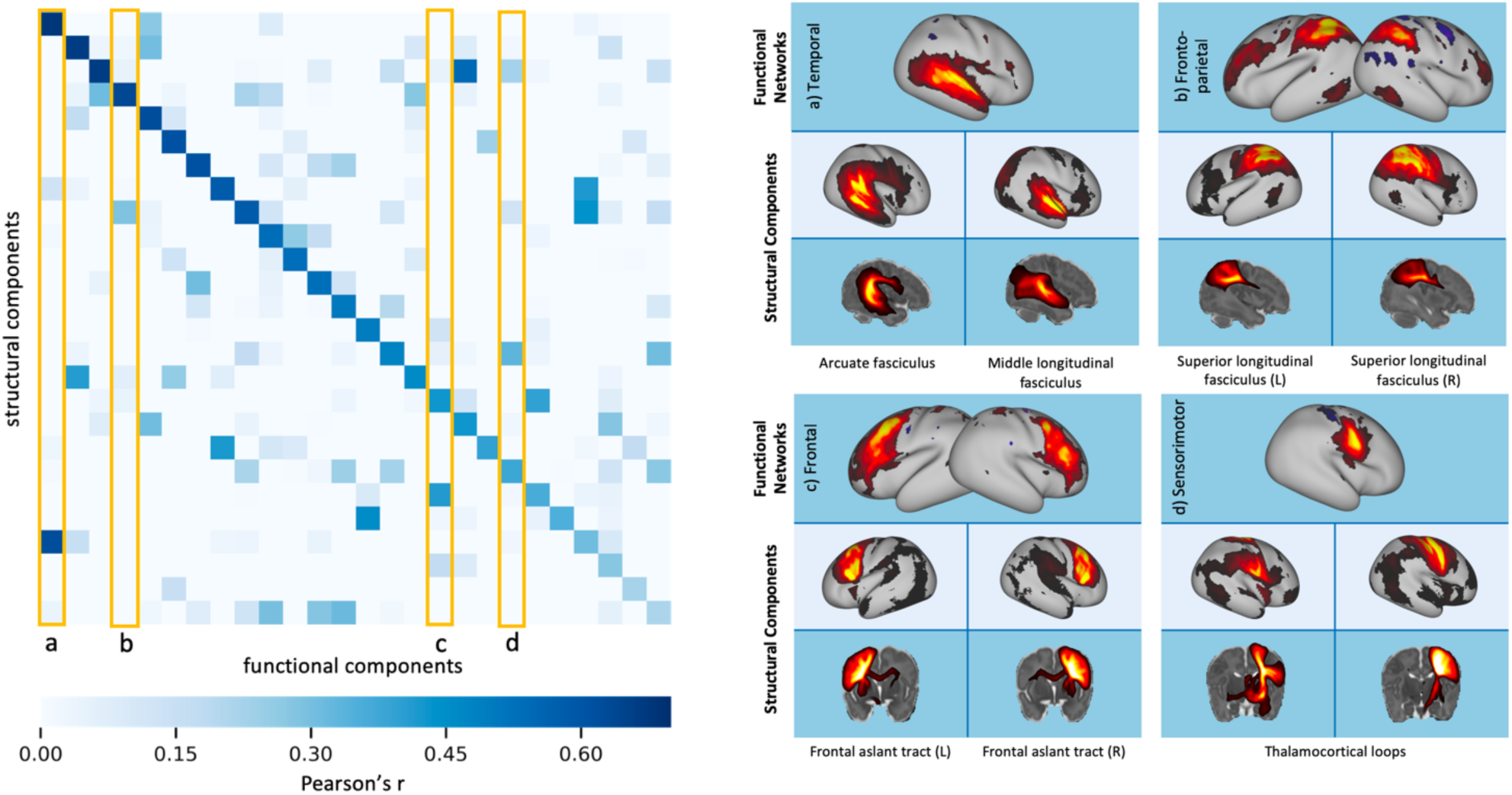
Left: correlation matrix between the fMRI RSNs and their 26 best-matched tractography NMF components. Right: examples of the functional networks and their most spatially similar grey matter components from structural NMF. These correspond to the columns outlined in yellow on the correlation matrix. The corresponding white matter patterns are shown as maximum intensity projects.

### Parcellations

The grey matter components from NMF were used to generate hard parcellations of the cortex, using a winner-takes-all approach. This process was carried out on each of the split-half groups to assess how robust the parcellations are to different groups of subjects. Figure 11 illustrates the parcellation results for different values of *K*. We can observe high reproducibility of the parcels between the two split-halves, and parcellation schemes are robust across different model orders. Figure 12a quantifies the similarity by showing the distributions of Dice scores across all generated parcels. This can be compared against distributions of Dice scores obtained from 100 random Voronoi parcellations (with spatial continuity-enforced) of the same order as the decomposition used in each case. The parcellations using the NMF components are significantly more consistent than the equivalent randomly generated parcellations. Also shown in figure 11, is a subject-specific parcellation generated from the results of a non-negative dual regression.

**Figure 11.**
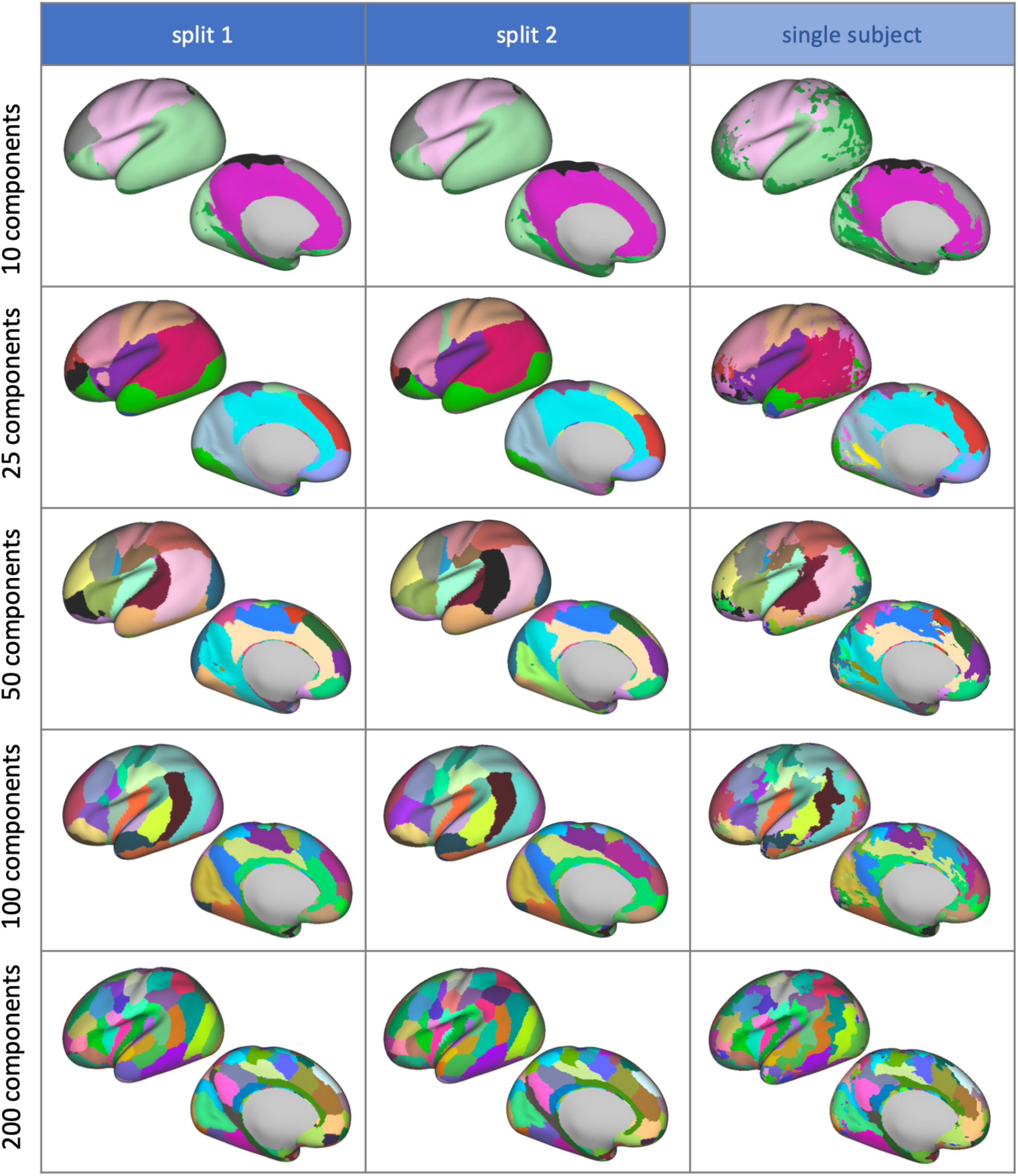
Hard parcellations of the cortical surface from NMF, from each split-half of the cohort and from dual regression of the group-level results onto a single subject. The left hemisphere displayed only. Parcels are colour matched according to the correlation values between the original grey matter components.

**Figure 12.**
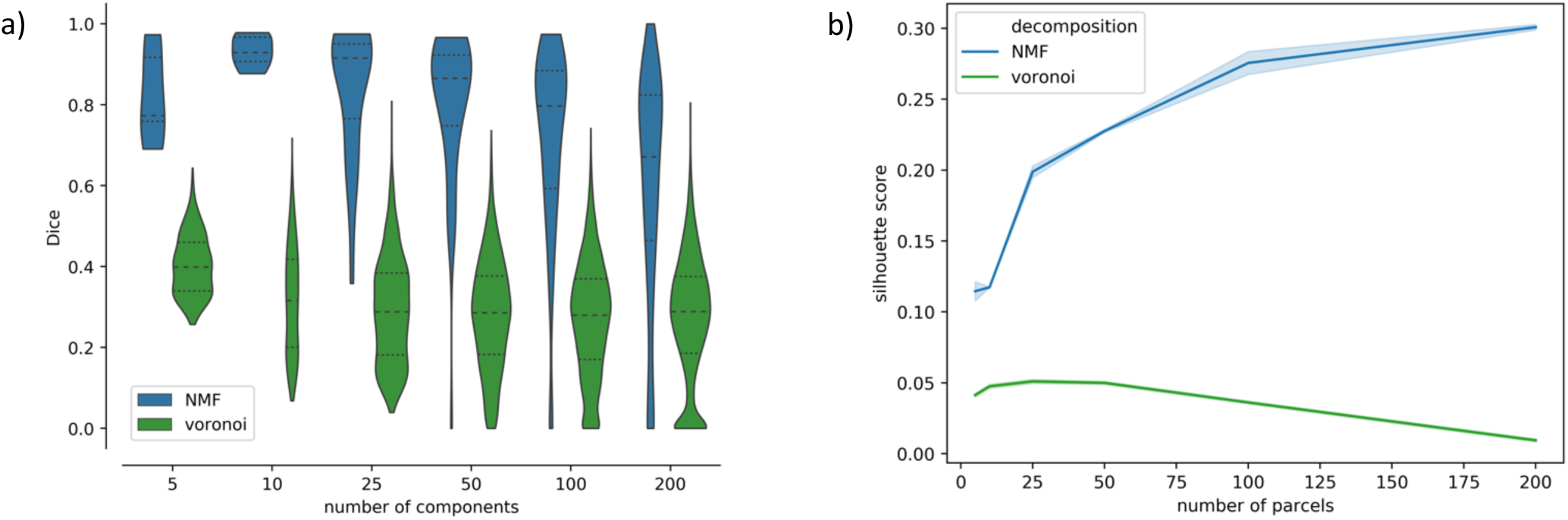
a) Dice scores of matching parcels across the split-half analysis. For comparison, we also calculated the Dice score between one of the splits’ NMF parcellations and 100 randomly generated Voronoi parcellations of the same model order. b) Mean Silhouette score across clusters for NMF and Voronoi parcellations with model orders of 5, 10, 25, 50, 100 and 200.

To further gain insight into the validity of these parcellations, we calculated the mean Silhouette score across parcels for the NMF-based parcellations at each model order, and for each split-half of the cohort. For comparison, we also computed the measure for 100 randomly generated Voronoi parcellations with the same number of parcels. A silhouette score measures the similarity of the data within a parcel, relative to their dissimilarity to data in other parcels. From figure 12b, we can see that the mean Silhouette score across parcels for our data-driven parcellations is consistently higher than for the equivalent random parcellations. Furthermore, we can see that the validity of the parcellations increases with increasing numbers of parcels in data-driven parcellations. On the contrary, for random parcellations, the Silhouette score peaks at *K* = 25, and then decreases for greater values of *K*. Our results show that our data-driven parcellations provide a more meaningful clustering of the data than random parcellations, even when the random parcellation has spatial contiguity enforced.

## Discussion

We have developed and demonstrated a non-negative framework for simultaneously mapping white matter connections and corresponding grey-matter networks from diffusion MRI data in a data-driven manner. We presented this approach within the context of mapping structural connectivity in the neonatal brain. Non-negative matrix factorisation (NMF) is a powerful alternative to traditional tract delineation that has no parametric assumptions, no dependence on predefined ROIs and masks in a template space, and is suited to the inherently non-negative nature of tractography data. We directly evaluated the performance of the framework using numerical simulated scenarios and indirectly explored the validity of the extracted components by comparing them against known tracts and against networks obtained from a different modality (resting-state fMRI). We also demonstrated benefits compared to a similar-in-spirit approach that used independent component analysis (ICA) to map connections in the adult brain (O’Muircheartaigh and Jbabdi, 2017). NMF is an alternative decomposition method that provides more interpretable and accurate reconstructions of non-negative sources than ICA.

Our work falls within the family of other data-driven approaches for mapping structural connections from whole-brain tractograms, such as (Garyfallidis et al., 2012; O’Donnell and Westin, 2007; Siless et al., 2018). Our approach extends these efforts by allowing simultaneous reconstructions of white matter bundles, but also corresponding grey-matter networks that these bundles connect. Furthermore, none of the previous data-driven approaches have been applied for mapping connections from diffusion MRI data of the neonatal brain, as shown here.

### Validation using Simulations

We used simulations to investigate the behaviour of the decompositions in controlled scenarios, in which the ground truth was known, and we could evaluate performance as a function of preselected features. In order to generate realistic simulations for such a decomposition framework, we therefore learned properties of the sources from distributions obtained from *in vivo* data, and mixed non-negative sources to generate synthetic data with a known number of components.

We first looked at the effect of adding an *L*_*1*_-norm regularisation term to the objective function for NMF (see equation 1). Increasing the regularisation reduces the accuracy of the data reconstruction, but a small amount (α = 0.1) improves the correlations between the sources and the components at lower model orders and promotes component sparsity. We decided to use an alpha value of 0.1 for subsequent work, as we deemed this to be a good compromise between higher component sparsity and sources reproduction, with only a minimal impact on reconstruction accuracy. Increasing the sparsity of components has been shown to generate features that are inherently more independent, while constraining the NMF solution space to make the decomposition more reliable (Hoyer, 2004).

We also looked at the effect of adding varying levels of Gaussian noise to the data. As expected, the reconstruction error of the decompositions increased with increasing noise, but the correlation between components and true sources was fairly stable, particularly at low model orders. Comparing the results from ICA and NMF, both were able to reconstruct the original data (using the dot product of the mixing matrix and component matrix) with good accuracy, but the components from ICA were less well matched to the true non-negative sources themselves than those from NMF. This is because the components from ICA contain negative values that are not found in the real sources, although mutual cancellation of positive and negative values in the components and mixing matrix allows the data matrix to be reconstructed accurately.

### Indirect Validation

White matter spatial maps of the NMF components show strong spatial similarity to known white matter pathways (Figures 6, 7, Suppl. Figure 4). We explored a range of model orders from 5 to 200. The lower model orders generate more distributed components that contain multiple white matter bundles, whereas the higher model orders give more specificity, as shown in Suppl. Figure 6. The components from lower model orders (eg. *K* = 5) are split into smaller constituent parts for higher model orders, providing a component hierarchy as *K* increases. Quantitative analysis of the components shows that reconstruction error decreases with more components and that the sparsity of the components increases. This reflects the higher degree of freedom afforded by more components that permit a more detailed reconstruction of the original data, and components that are more tightly localised around fine-grained regions of similar connectivity. NMF components are more sparse than those from ICA, which indicates that the former is able to localise connectivity patterns more effectively, disregarding redundant information and keeping non-negativity in the reconstruction. At the same time, putting sparsity and sign aside, the patterns from ICA and NMF look broadly similar, as seen in figure 6. This hints towards NMF being able to separate spatially independent components, in an analogous manner to ICA, despite not having independence constraints enforced directly. This is because the sparsity constraint on the NMF decomposition promotes non-Gaussianity in the resultant components, which is used as a proxy for independence in the FastICA algorithm (Hyvärinen and Oja, 2000). Indeed, sparsity and independence criteria have previously been shown to generate very similar basis sets across several different data types (Saito et al., 2000).

The grey matter maps of the NMF components were also shown to align well to resting-state networks from fMRI. This provides further evidence that these data-driven results are anatomically meaningful. It also opens up future possibilities for devising a multi-modal data-driven framework that can fuse information across modalities and perform decompositions simultaneously for dMRI and fMRI data.

### Parcellations

We used the grey matter maps of the NMF components to generate a cortical parcellation scheme. Specifically, each vertex on the cortical mesh was labelled according to the component with the strongest weighting at each point. This leads to a parcellation in which clusters share similar patterns of structural connectivity to the rest of the brain. Depending on the model order of the decomposition, the parcellation can be coarse or more fine-grained (see figure 11). An advantage of this approach is that it is entirely data-driven, so the parcellations are not biased by any subjective measures. It can also be used to generate subject specific parcellations, by using the subject-level grey-matter maps from dual regression. Other cortical parcellation schemes that exist for neonates tend to be based on adult atlases (Alexander et al., 2017; Oishi et al., 2011). Cortical parcellation schemes based on structural connectivity exist for adults, see (Tittgemeyer et al., 2018) for a recent review, but not for babies.

We also performed a split-half reliability analysis of the parcellations, using Dice Score as a similarity measure, to see how reproducible the parcellations are for different model orders. We compared the results with the Dice score between one split and a set of randomly generated Voronoi parcellations. For all model orders, the data-driven parcellations were more consistent than random parcellations. In addition, we used Silhouette score as a measure of the parcel validity, and again compared the performance of the NMF-based parcellations against 100 random Voronoi parcellations. Silhouette score measures the similarity of the connectivity profile of a given grey matter vertex to others in its parcel, relative to the connectivity of vertices in other parcels. We found that our data-driven parcellations consistently scored higher on this measure than the random parcellations (see figure 12). However, although this demonstrates that our parcellations are effectively clustering similarly connected vertices, this does not necessarily indicate that we are delineating neurobiologically relevant areas, which can be heterogeneous in their connections (Van Essen and Glasser, 2018). Incorporating information from different modalities would help to improve these parcellations further and ensure that the areal boundaries are neurobiologically relevant (Glasser et al., 2016). NMF can assist in enabling such benefits from multi-modal semi-automated approaches.

### Decomposition Domain

In the results presented here, we have been applying decompositions in the WGB seed domain, allowing white matter tract overlap. We also tried applying the decompositions to the transpose of the connectivity matrix, **X**^**T**^, which meant decomposing (and in the case of ICA enforcing independence) in the tract domain. ICA and NMF were performed on the transpose of the split 1 connectivity matrix, with *K* = 50. Looking at the similarity between the results from both methods (see Suppl. Figure 8), we can see that the ICA components are most affected by this change. Most NMF components are nearly identical to the original results. This agrees with expectations, as in NMF the sparsity and non-negativity constraints are enforced in both the mixing matrix and the components (see equation 1).

### Limitations

Our decomposition framework uses whole-brain tractography data and its performance can therefore be challenged by tractography limitations, which are important to keep in mind when interpreting results. Tractography is an indirect measure of anatomy that is prone to identifying false positive connections (Maier-Hein et al., 2017). It has also been shown that tractography streamlines are biased towards terminations in the gyri rather than the sulci (Schilling et al., 2018; Van Essen et al., 2013), although the effects of this “gyral bias” can be minimised by seeding from the cortical surface rather than the whole brain (Donahue et al., 2016; Schilling et al., 2018), as we have done here. We have also shown in previous work that the effects of gyral bias are less prevalent in neonates than in adults due to the less developed cortical folding (Thompson et al., 2019) and we therefore expect less direct influence of such biases into the NMF performance in the neonatal brain. In fact, our parcellation borders did not show a consistent overlap with sulcal fundi or gyral crowns (Suppl. Figure 9).

Data-driven decompositions can be more computationally demanding than standard tractography approaches, as they consider all data at once and extract all white-matter and grey-matter maps simultaneously, within the same decomposition. To reduce the memory requirements and the computational burden, we binned the whole brain tractography data into a 2 mm spatial grid, which subsequently defined size *M* in the decompositions (Figure 1); rather than using the native 1.5 mm spatial grid of the dMRI data. This provides WM components at a lower resolution than available in the original data but does not change any trends or conclusions drawn from the presented analyses.

## Conclusions

We have shown that data-driven methods can be used to jointly map white matter bundles and their corresponding grey matter networks from dMRI tractography data from neonatal subjects. In particular, we show that non-negative matrix factorisation provides a robust decomposition that is a natural fit for the inherently non-negative structural connectivity data.

## Acknowledgements

E.T. is supported by funding from the Engineering and Physical Sciences Research Council (EPSRC) and Medical Research Council (MRC) [grant number EP/L016052/1]. S.N.S. is also supported by grant [217266/Z/19/Z] from the Wellcome Trust. Data were provided by the developing Human Connectome Project, a KCL-Imperial-Oxford Consortium funded by the European Research Council under the European Union Seventh Framework Programme (FP/2007-2013) / ERC Grant Agreement no. [319456]. We are grateful to the families who generously supported this trial. The computations described in this paper were performed using the University of Nottingham’s Augusta HPC service and the Precision Imaging Beacon Cluster, which provide High Performance Computing service to the University’s research community.

## Supplementary Figures

**Supplementary Figure 1.**
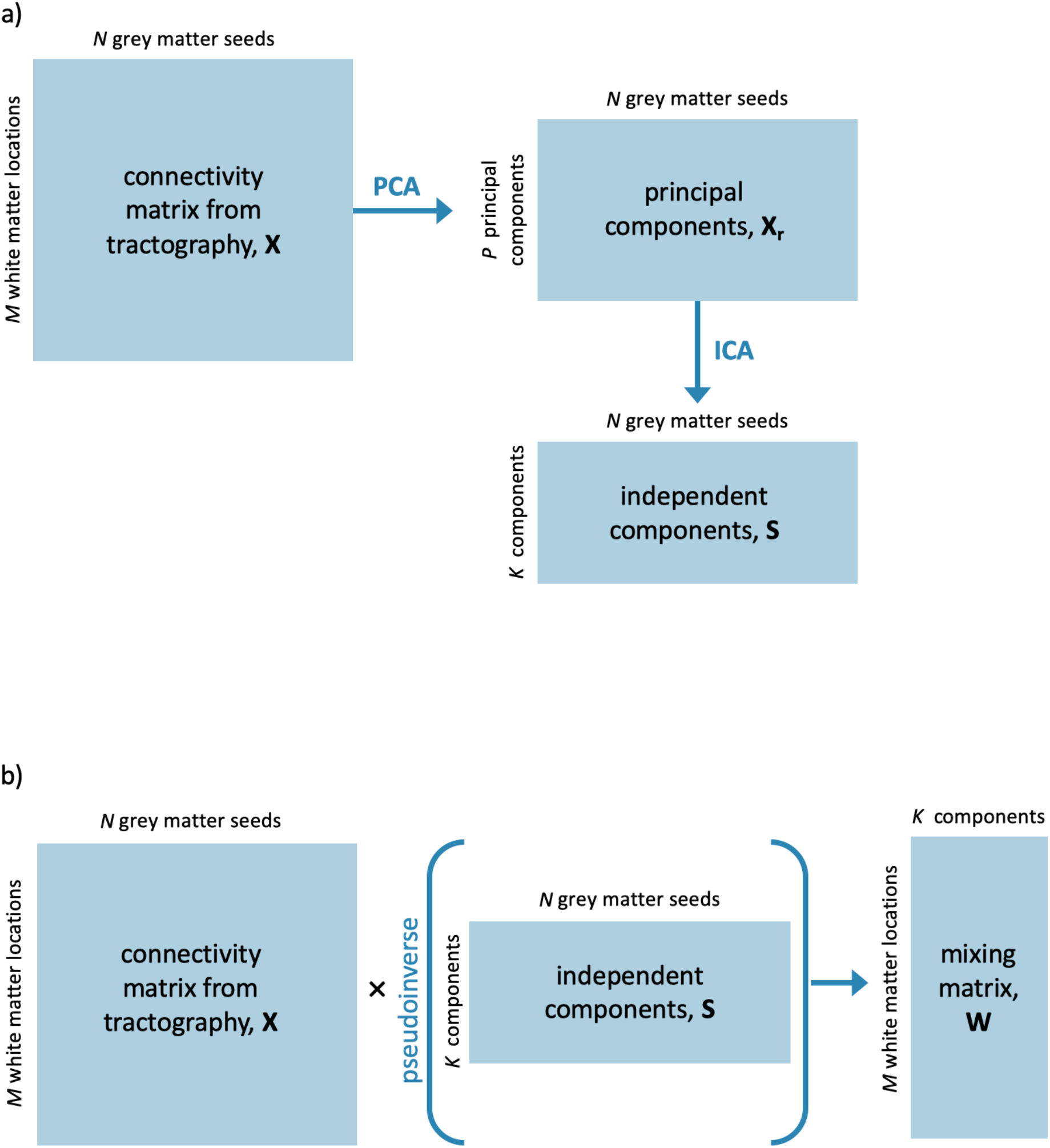
Illustration of a back projection step to obtain a white matter mixing matrix after data reduction by PCA. a) Data are first reduced to P principal components, and ICA applied to the data in the reduced subspace. b) Independent components are regressed onto the original data to obtain a mixing matrix in white matter space.

**Supplementary Figure 2.**
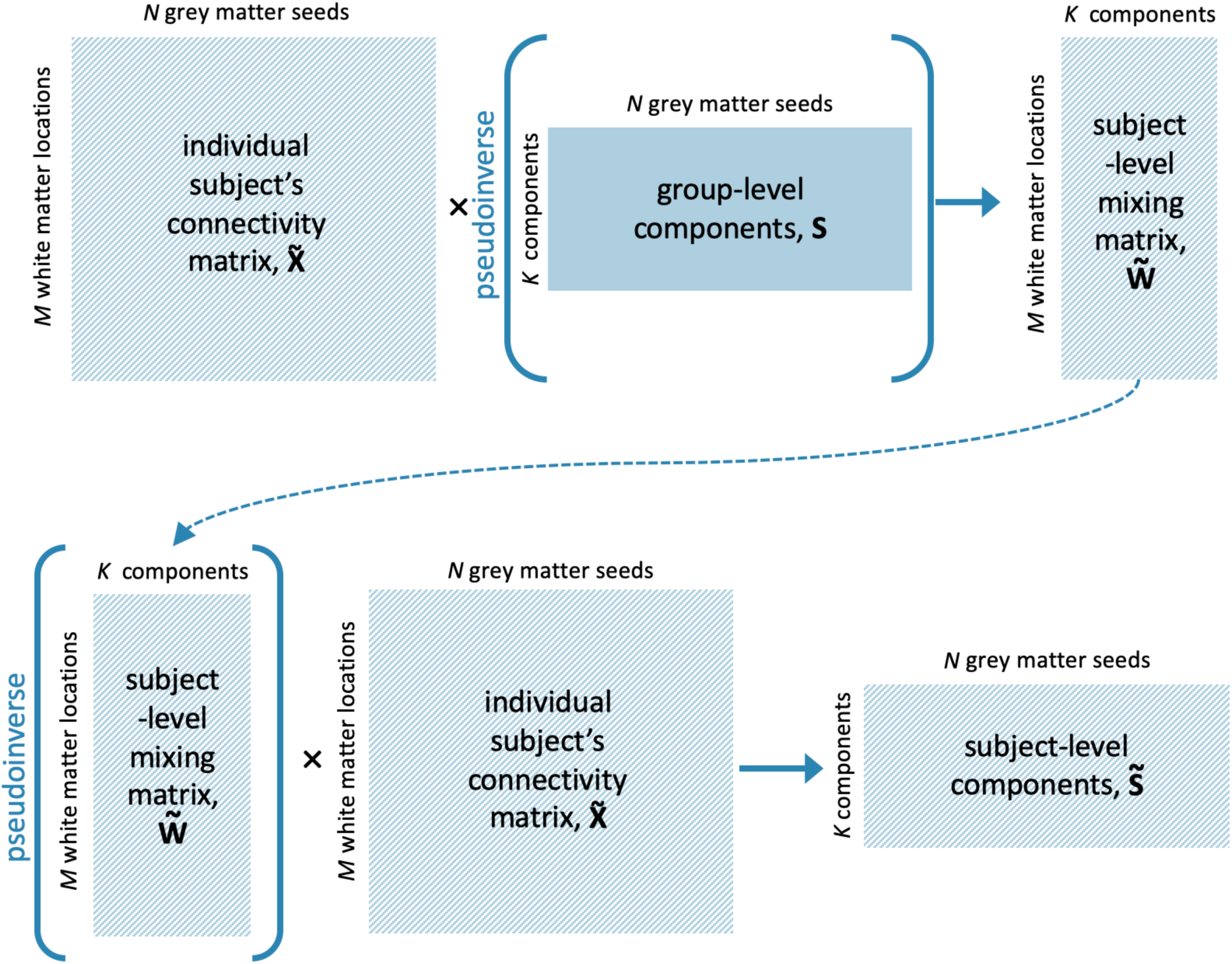
Schematic illustration of the dual regression step used for ICA, that generates subject-level representations of the group-level components. The group-level grey matter components are first regressed onto the subject’s connectivity matrix to obtain the subject-level representations of the components in white matter. We then use the pseudoinverse of this mixing matrix to obtain the subject-level grey matter components.

**Supplementary Figure 3.**
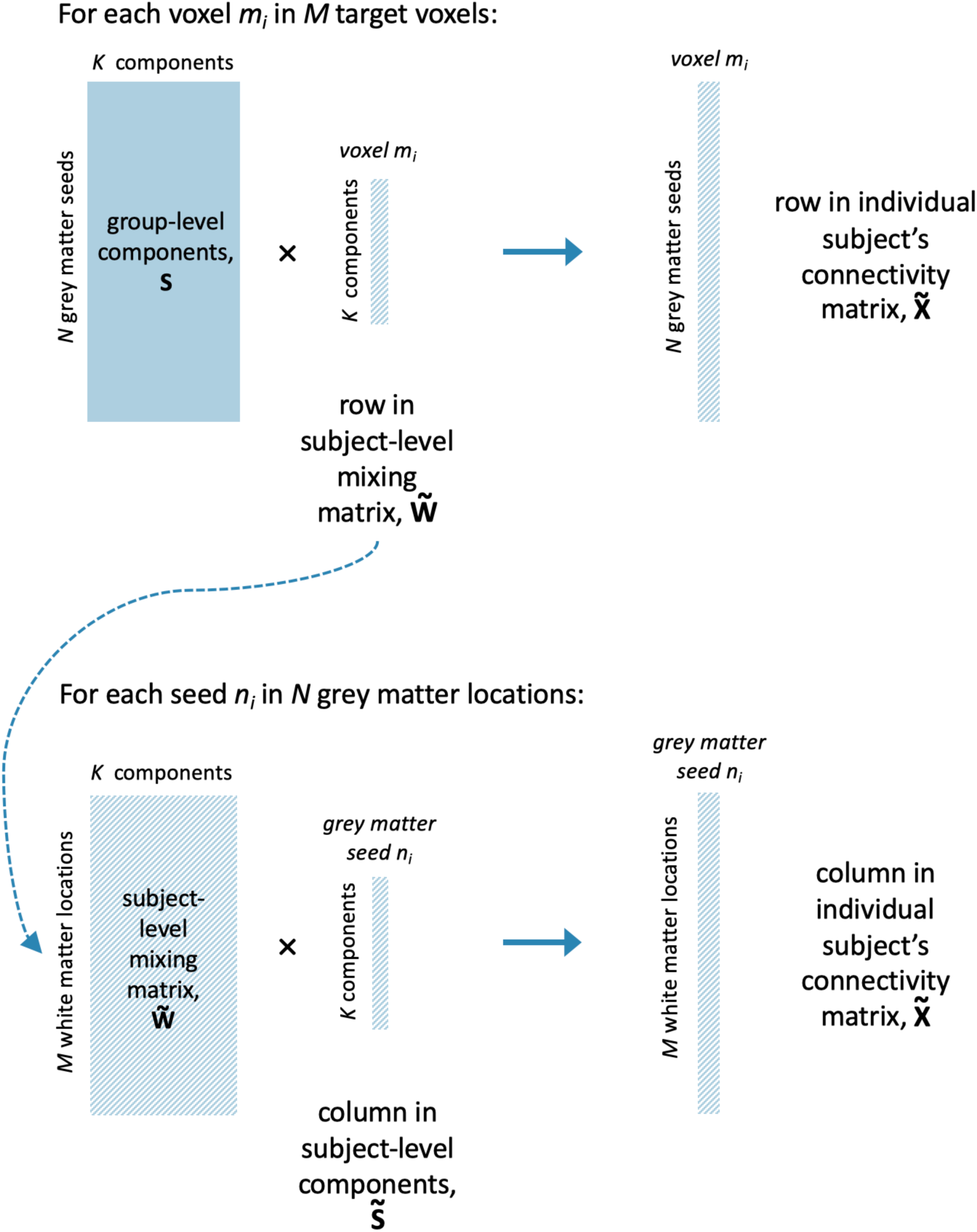
Schematic illustration of the non-negative dual regression step used for NMF, that generates subject-level representations of the group-level components. We first use NNLS to solve the top equation, using the group-level grey matter components and the subject’s connectivity matrix to solve for each row in the subject-level mixing matrix. We then use this mixing matrix to find the subject-level grey matter components by the same method.

**Supplementary Figure 4.**
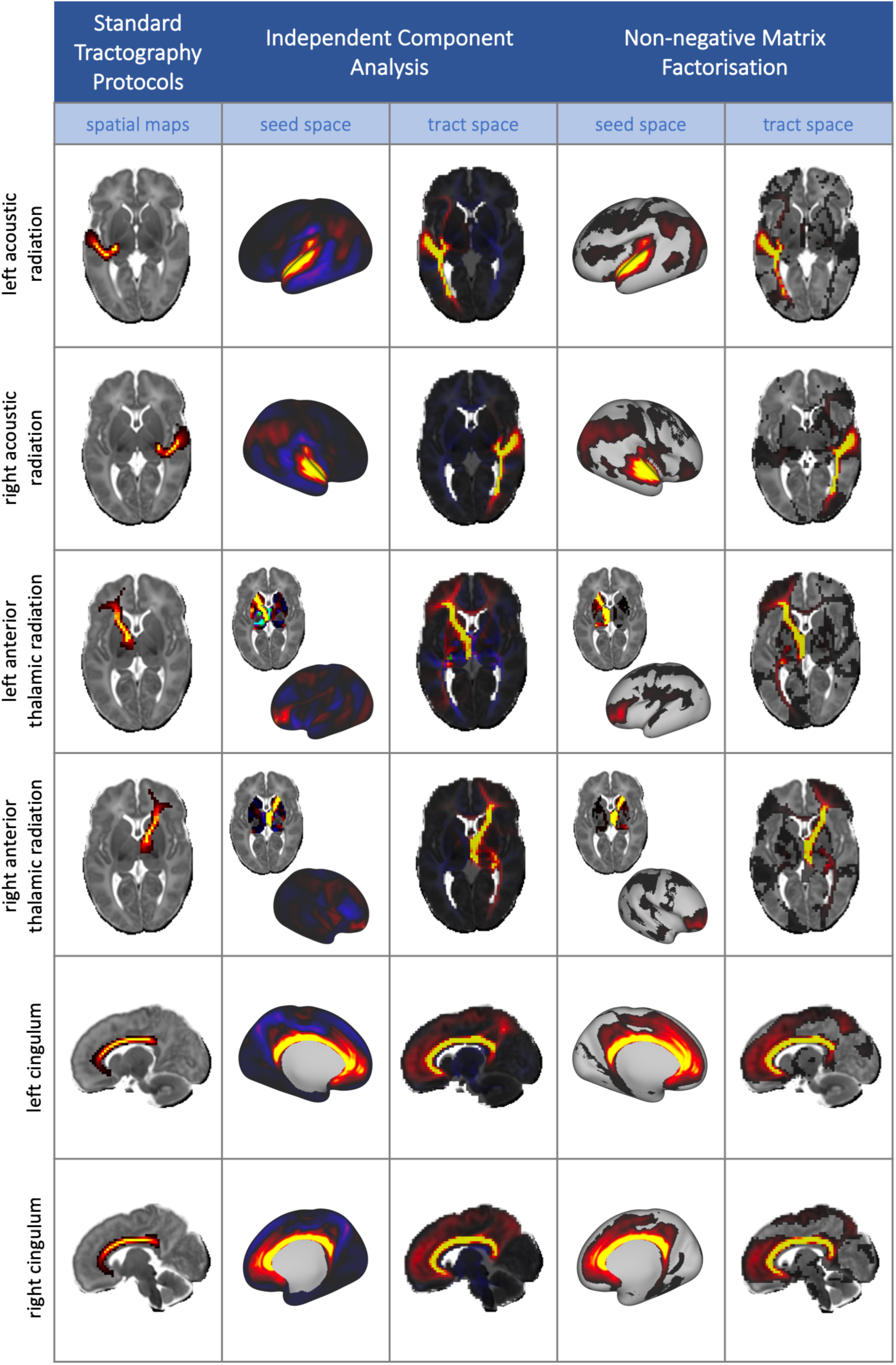
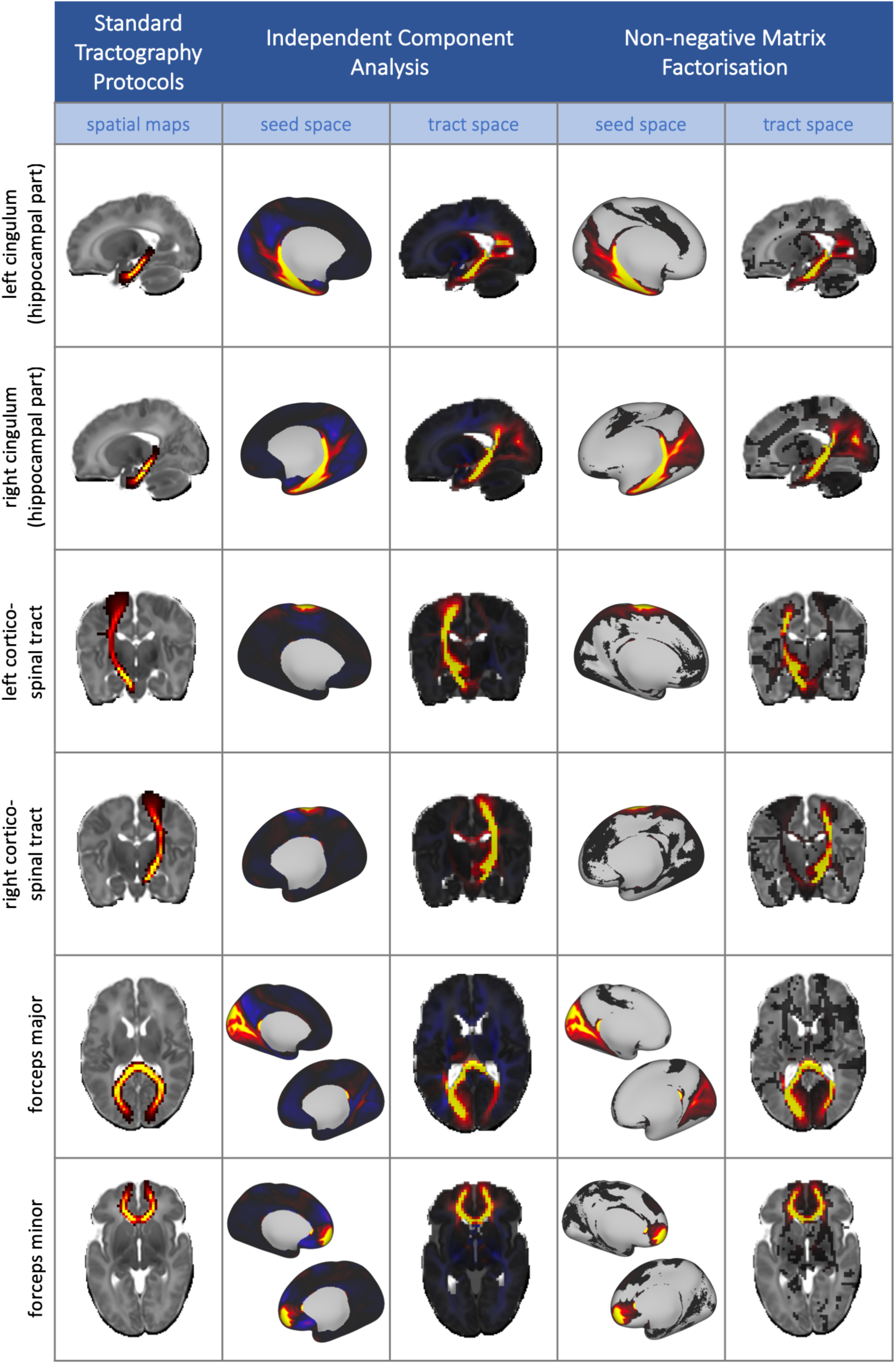
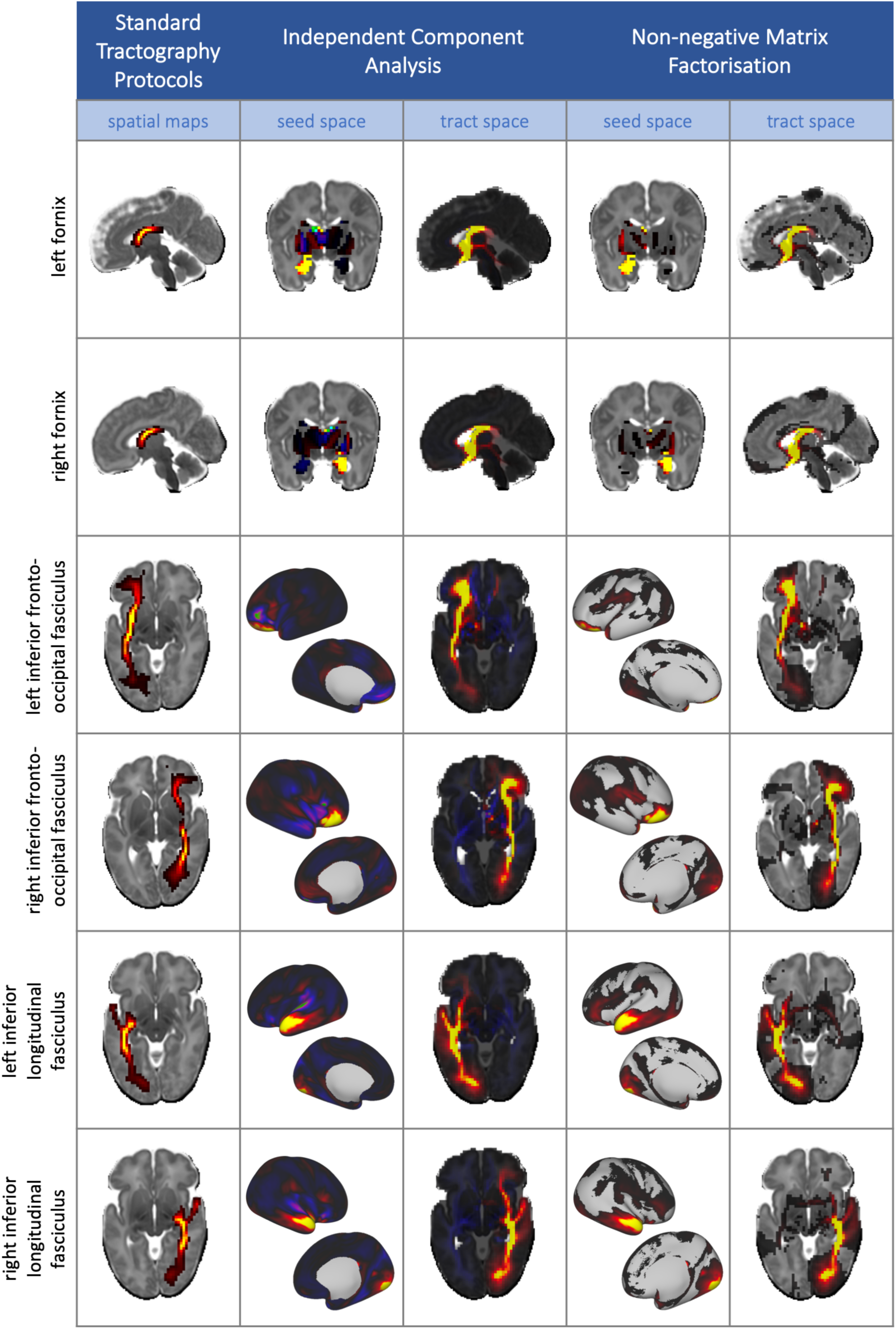
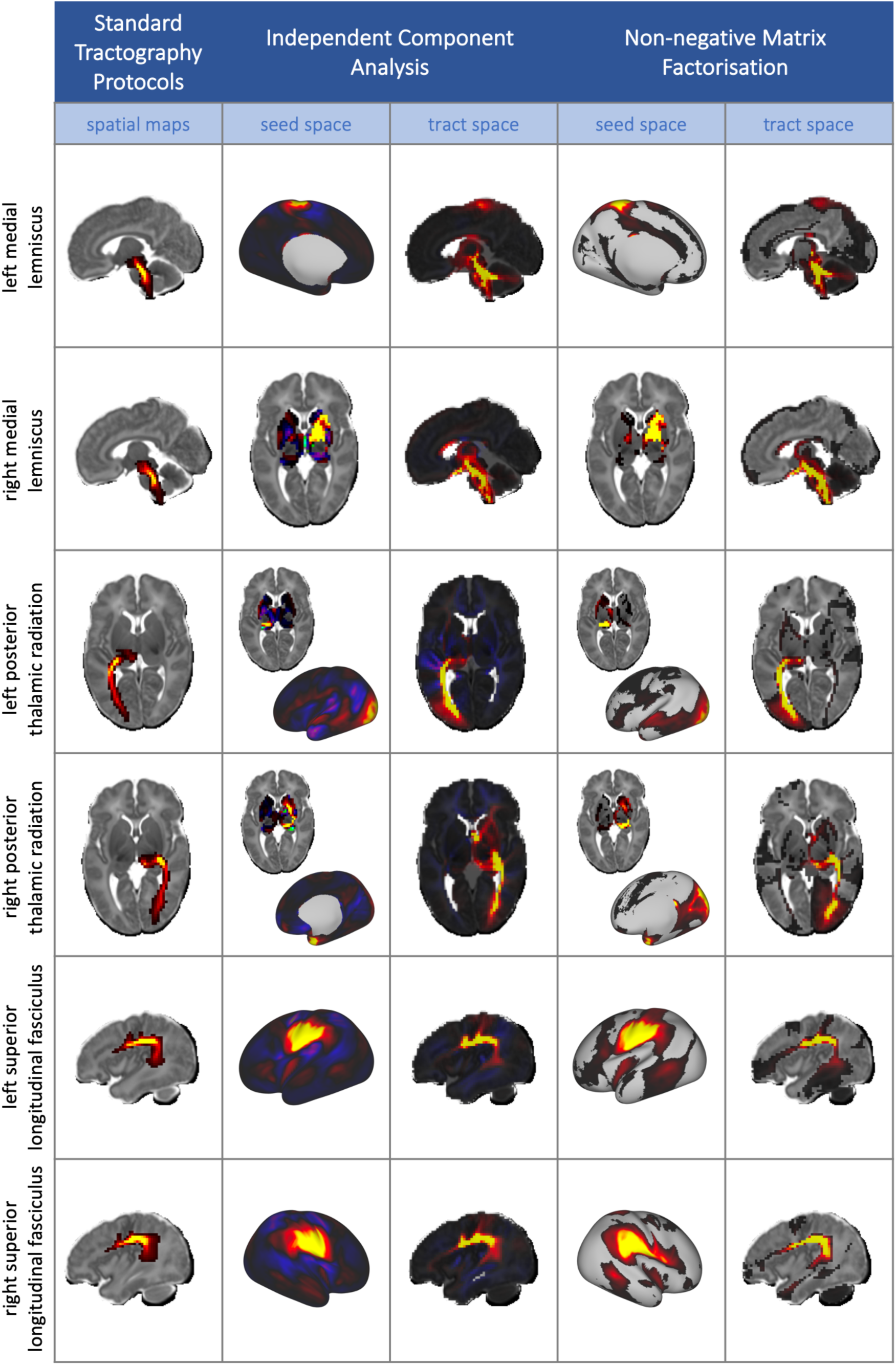
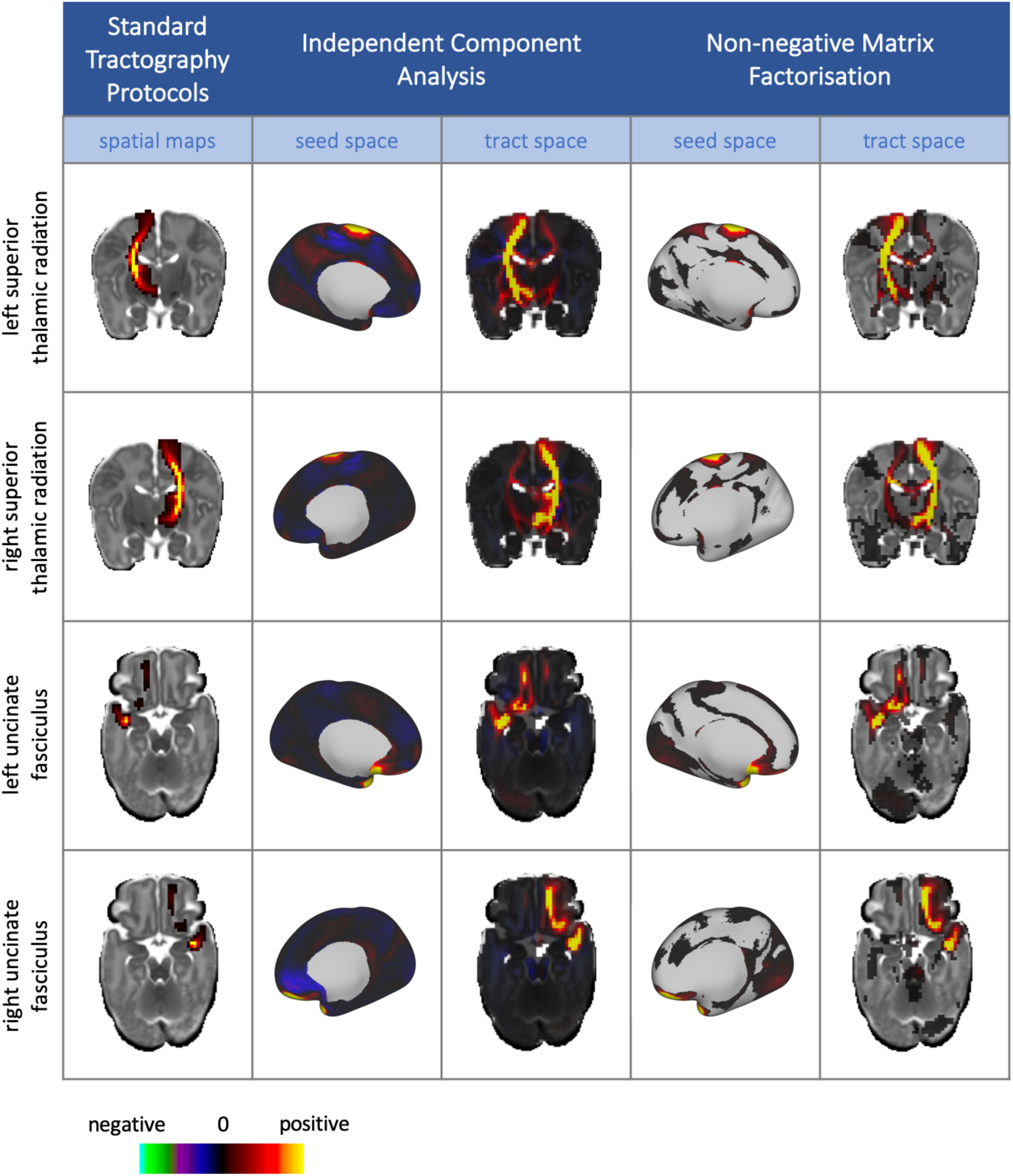
The full set of 28 tracts from the standardised protocols (Bastiani et al., 2019), alongside there corresponding components from ICA and NMF. Data-driven components are unthresholded to enable the comparison between the negative values in the ICA components and the sparse, non-negative representations from NMF. All tractography and data-driven results are taken from split 1 of the split-half analysis.

**Supplementary Figure 5.**
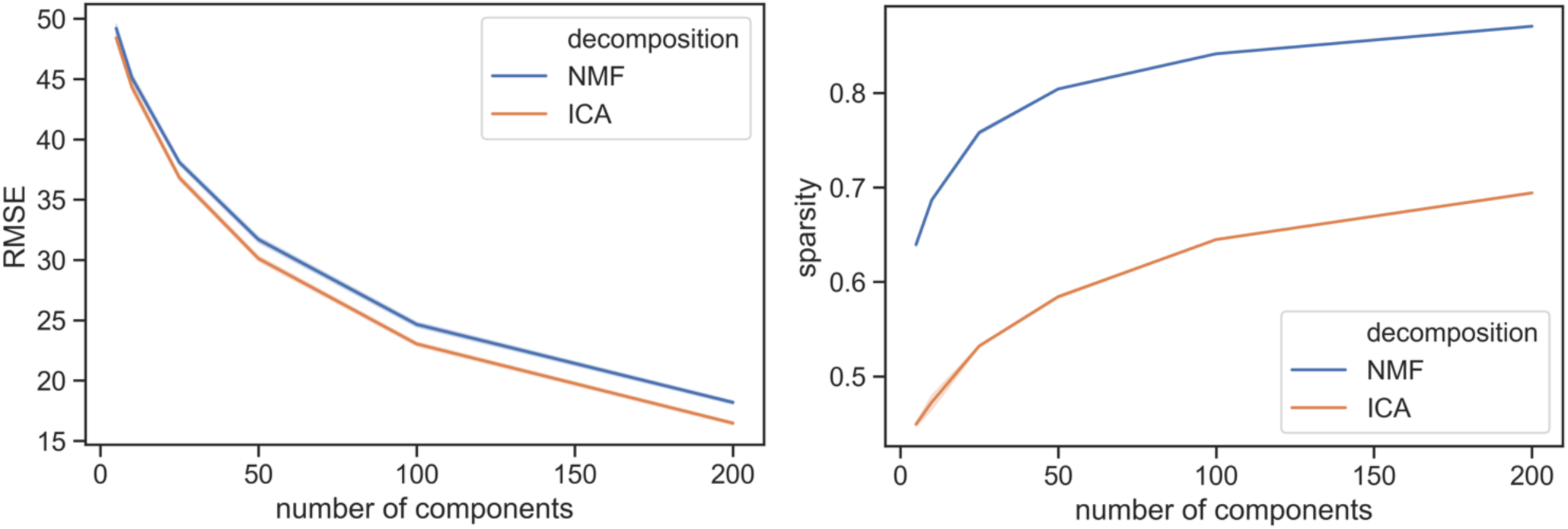
a) Reconstruction error at each model order for ICA and NMF. This is the root mean squared error between the connectivity matrix and the dot product of the mixing matrix and the components. b) The sparsity of the derived components, calculated according to equation (2).

**Supplementary Figure 6.**
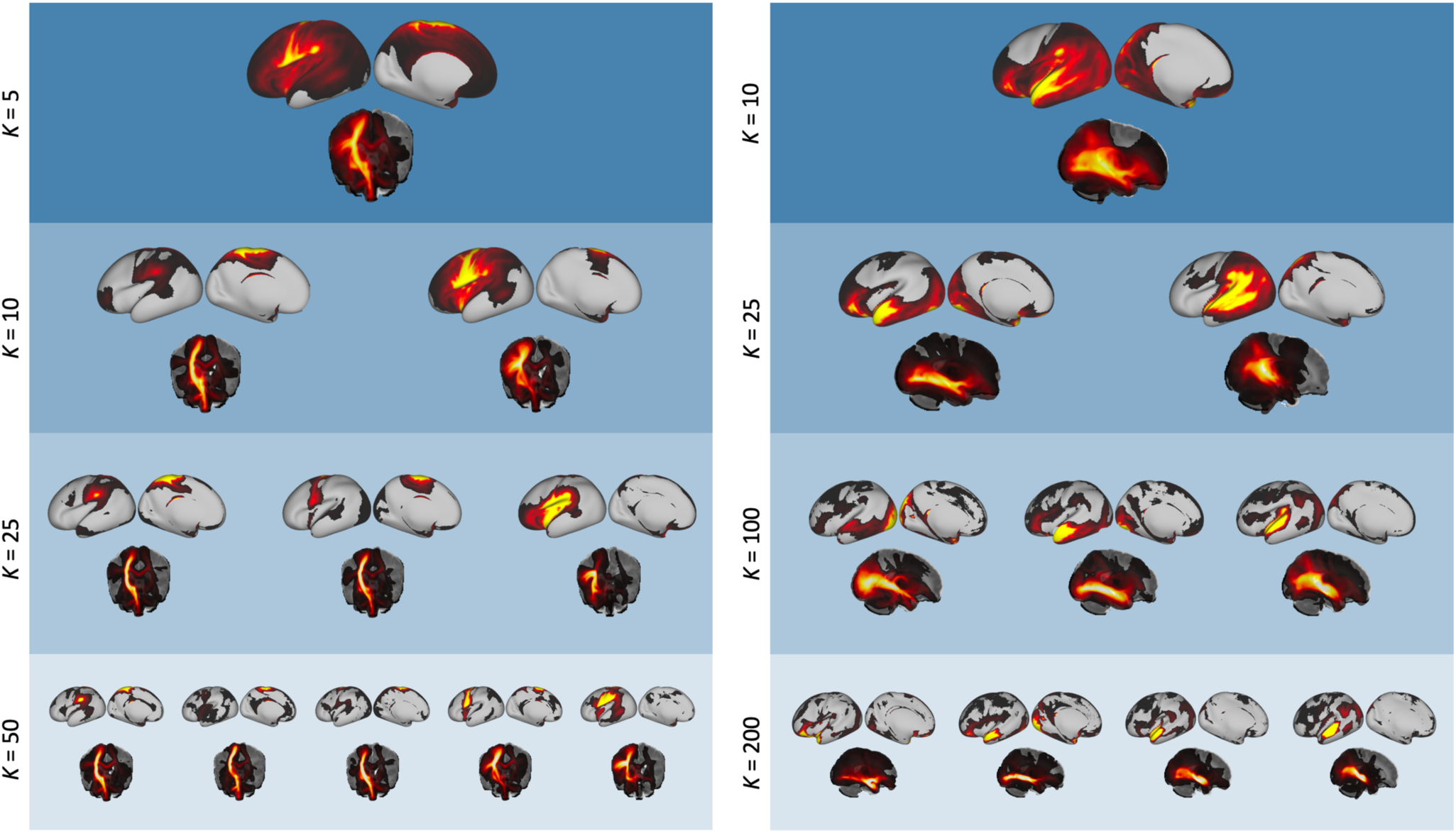
Similar components across different model orders, demonstrating the hierarchical nature of the decomposition. Starting with a single component from a lower dimensionality decomposition, we show components from higher model orders that have high spatial correlation with the original component, in tract space (r > 0.5). Tract space results are displayed as maximum intensity projections. Left: A component from the K = 5 decomposition showing the left cortico-spinal tract, which is split into more localised sub-components for higher K. Right: A component from the K = 10 decomposition that includes several different association fibres in the left hemisphere. At K = 200, this has been split into the uncinate fasciculus, inferior longitudinal fasciculus and middle longitudinal fasciculus.

**Supplementary Figure 7.**
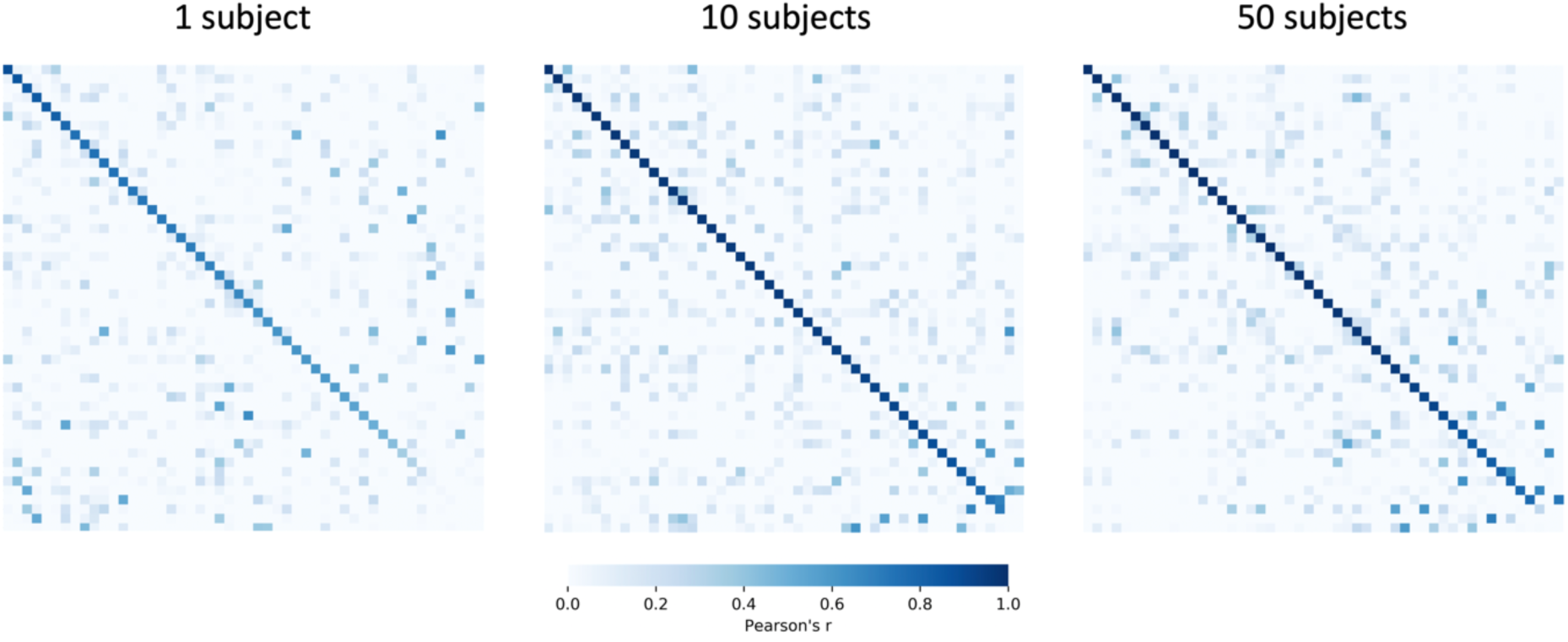
Correlation matrices between the white matter K = 50 NMF spatial maps from smaller group-level decompositions with the split 1 components from the main analysis. Each entry corresponds to the correlation between a component obtained from a group of L subjects (L=1,10,50) and the best-matching component from one of the split-halves of the full cohort (161 subjects). The matrices have been reordered so that matching components lie on the diagonal and are in descending order in terms of correlations. We can see that there is no significant increase in agreement between the 10 subject group with the full cohort and the 50 subject group and the full cohort (average Pearson’s r of matching components = 0.89 for both), which indicates that the method is robust even for small group sizes.

**Supplementary Figure 8.**
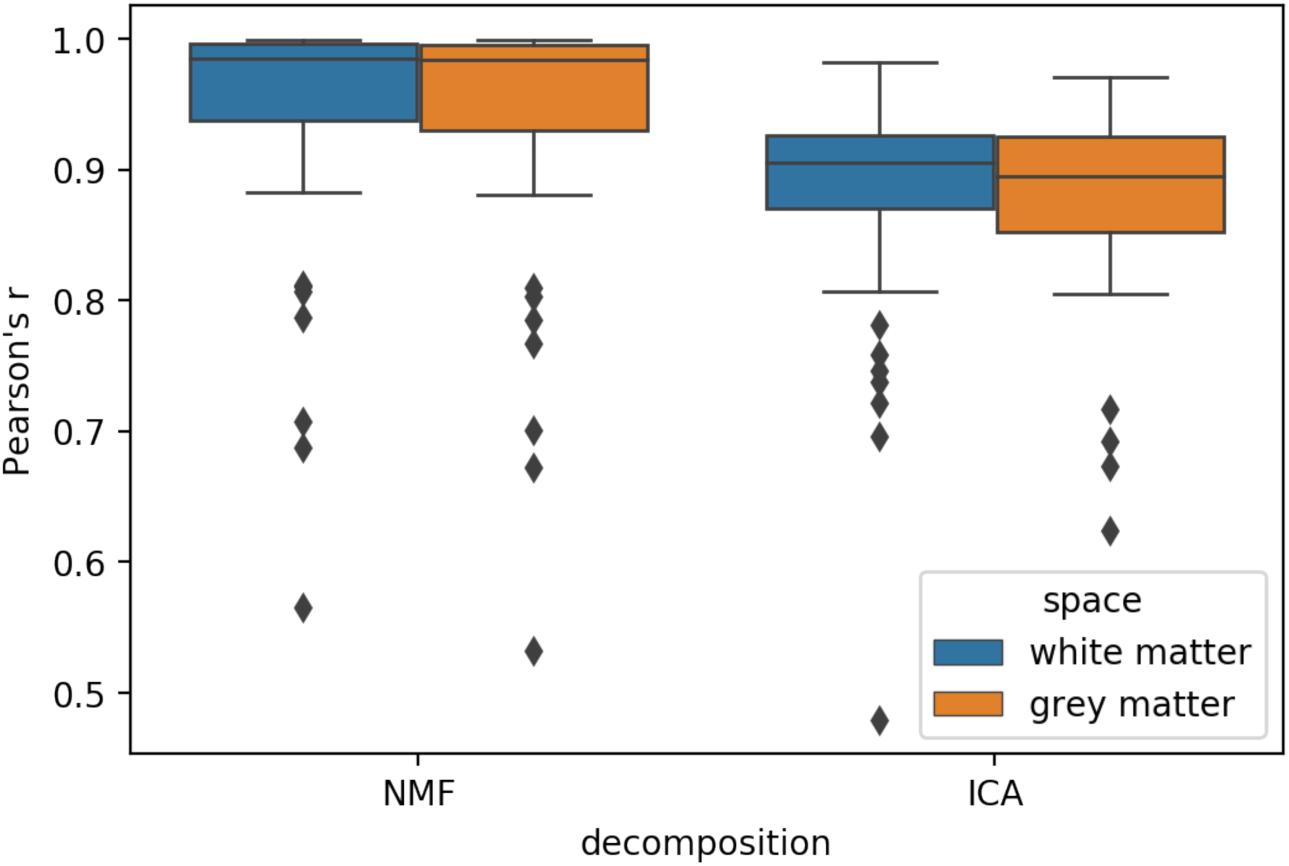
Correlations between decompositions applied to connectivity matrices in both the (seed x white matter) and (white matter x seed) configuration. Highest Pearson’s r is plotted for each component or column of the mixing matrix with the equivalent matrix from the transposed decomposition.

**Supplementary Figure 9.**
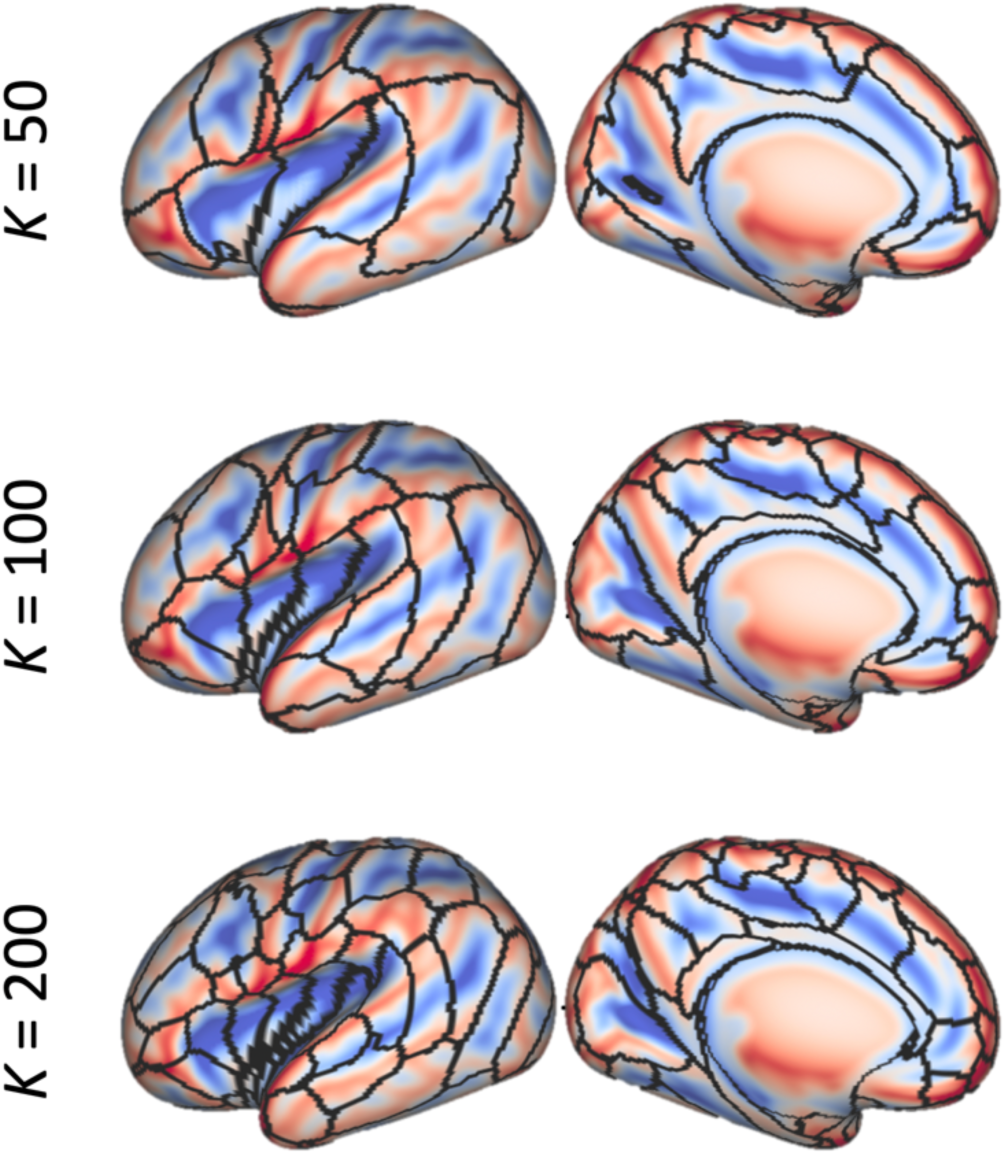
Our group-level NMF parcellation overlaid on the dHCP’s 40-week PMA sulcal depth template (Bozek et al., 2018) (blue for sulcal fundi, red for gyral crowns). There is no consistent pattern for parcellation boundaries to follow sulci or gyri, which indicates that our parcellation is not driven (at least to a large degree) by the gyral bias.

## Footnotes

1 By “dense” we refer to voxel-wise / vertex-wise representations rather than areal-wise nodes, i.e. N and M are in the order of thousands.

2 www.developingconnectome.org/second-data-release

